# Weighted 2D-kernel density estimations provide a new probabilistic measure for epigenetic age

**DOI:** 10.1101/2024.06.10.598169

**Authors:** Juan-Felipe Perez-Correa, Thomas Stiehl, Riccardo E. Marioni, Janie Corley, Simon R. Cox, Ivan G. Costa, Wolfgang Wagner

## Abstract

**Background:** Epigenetic aging signatures can provide insights into the human aging process. Within the last decade many alternative epigenetic clocks have been described, which are typically based on linear regression analysis of DNA methylation at multiple CG dinucleotides (CpGs). However, this approach assumes that the epigenetic modifications follow either a continuous linear or logarithmic trajectory. In this study, we explored an alternative non-parametric approach using 2D-kernel density estimation (KDE) to determine epigenetic age.

**Results:** We used Illumina BeadChip profiles of blood samples of various studies, exemplarily selected the 27 CpGs with highest linear correlation with chronological age (R^2^ > 0.7), and computed KDEs for each of them. The probability profiles for individual KDEs were further integrated by a genetic algorithm to assign an optimal weight to each CpG. Our weighted 2D-kernel density estimation model (WKDE) facilitated age-predictions with similar correlation and precision (R^2^ = 0.81, median absolute error = 4 years) as other commonly used clocks. Furthermore, our approach provided a variation score, which reflects the inherent variation of age-related epigenetic changes at different CpG sites within a given sample. An increase of the variation score by one unit reduced the mortality risk by 9.2% (95% CI (0.8387, 0.9872), P <0.0160) in the Lothian Birth Cohort 1921 after adjusting for chronological age and sex.

**Conclusions:** We describe a new method using weighted 2D-kernel density estimation (WKDE) for accurate epigenetic age-predictions and to calculate variation scores, which provide an additional variable to estimate biological age.

## Background

Aging is reflected by gains and losses of DNA methylation (DNAm) at specific sites in our genome and this can be used to determine donor age. Epigenetic clocks have become a central component of aging research (1). Since they were first described (2, 3), epigenetic age-predictors have become more and more sophisticated and tailored toward specific applications (4). First generation clocks are trained to correlate as highly as possible with chronological age, e.g. for applications in forensics. Even in these clocks, the deviation of chronological and epigenetic age (delta age) is often indicative for all-cause mortality and affected by various diseases (5). However, this phenomenon has been proven to decrease when the training sample size is large enough or when a correction for the white blood cells counts is applied (6). To better reflect the individual pace of the aging process, multifactorial second generation clocks have been described. In addition to age, they also integrate epigenetic indicators of other clinical parameters, such as blood counts, glucose levels, or blood pressure. Furthermore, third generation clocks have been trained on large cohort studies and implement additional clinical measurements to better quantify the individual pace of aging (7).

Despite the wide range of epigenetic clocks that have already been established, most of them rely on the same approach: signatures of age-associated CG dinucleotides (CpGs) are preselected in a training dataset and then used for multivariable regressions under the assumption that there are linear changes of methylation with age. Particularly in childhood, age-associated DNAm changes follow a logarithmic pattern (8), which can be effectively addressed through non-linear corrections (9) – but the predictors still follow the assumption that there is a continuous trajectory – either linear or logarithmic. Recently, alternative non-linear epigenetic clocks have been described that are based on deep learning, e.g. by neuronal network frameworks in multidimensional space (10-12). Furthermore, machine learning algorithms can be used to derive non-parametric epigenetic clocks based on Gaussian process regression models (13). While these approaches theoretically use the full range of DNAm information of the training dataset, they are relatively complicated. Furthermore, all of these epigenetic clocks only provide one specific age-prediction for a given sample as a single output.

In this study, we introduce a novel methodology to build epigenetic clocks based on 2D kernel density estimation (KDE). For each individual CpG of the aging signature, the KDE puts age and DNAm into relation within a matrix of densities that is translated into probabilities. Integration of these probabilities can then be used to determine the most likely age-prediction. Our KDE approach is non-parametric and does not require linear or logarithmic assumptions. We demonstrate that, particularly with a weighted KDE model, we can facilitate robust epigenetic age-predictions. Furthermore, the probability distribution provides insight into how consistent the age-associated DNAm is for a specific age-prediction in different sites of the genome. This variation score is a new complementary measure that may be useful to estimate biological age.

## Methods

### DNA methylation data

We compiled DNAm datasets of human peripheral blood samples of 10 different studies from Gene Expression Omnibus (GEO; Supplemental table S1). We considered only samples that were classified as healthy or controls. Datasets were separated into a training set (7 Illumina HumanMethylation450K array studies, 1029 samples, age range 1-101 years, 50.1% female) and an independent validation set (3 Illumina HumanMethylation450K array studies, 628 samples, age range 2-79 years, 53.4% female). The data was processed in R 4.3.0. with the geoGEO function of the GEOquery package. CpGs in sex chromosomes and 6749 single nucleotide polymorphism (SNP) probes (according to the HumanMethylation450K v1.2 annotation) were filtered out.

### Epigenetic clock based on kernel density estimation

For selection of age-associated CpGs, we have exemplarily focused on the CpGs with highest linear correlation with chronological age in the training set: either 27 CpGs with R^2^ > 0.7 (Supplemental table S2), or 491 CpGs with R^2^ > 0.6 (Supplemental table S3). To generate KDE clocks based on these CpGs we tested various alternative approaches. Initially, we used all 1029 samples of the training set to generate 2D kernel density plots of chronological age *versus* DNAm. Since this approach resulted in offsets of epigenetic age-predictions due to the heterogeneous age-distribution in the training set, we subsequently adjusted densities of the original kernel by the frequency of donor ages. To this end, we only considered samples ≤ 85 years to avoid a distortion for age-categories with very few samples. Alternatively, we tested conditional density resampled estimate of mutual information (DREMI) (14), but this did not improve age-predictions (data not shown). Finally, we randomly selected 15 samples of the training set for each 5 year bin from 0 to 90 years, and for one additional bin with samples older than 90 years. This resulted in a subset of 285 samples with homogeneous age distribution.

For each CpG, the kernels were generated with the function kde2d of the MASS R package, with 101 grid points, resulting in kernels with age on the x-axis (0 – 100 years), beta values on the y-axis (0 to 100% DNAm) and the density on the z-axis. Thus, each CpG kernel corresponds to a 3D matrix *K*, where *K*_*i*,*j*,*k*_ represents the density value of the CpG *k* for a sample with age *i* and DNAm percentage *j*. At this point, a preliminary probabilistic kernel age prediction for a sample can be done by:

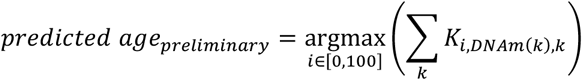

where *k* iterates through the selected CpGs and DNAm(*k*) is the DNAm percentage of the CpG *k* in the considered sample. For each CpG *k*, the row corresponding to the DNAm(*k*) is a vector of 101 densities, each of them corresponding to the years 0 to 100, respectively. After performing the summation of all these density vectors, the algorithm returns the age *i*, the one with the maximum cumulative density of the summation, as the final age.

To improve the performance of the predictor, a genetic algorithm was implemented to include a weight coefficient *w*_*k*_ for each CpG *k*, with possible values in the range [-10,10]. For this, the function *minimize* from the R package *EmiR* was used, with a total of 150 initial conditions, 100 iterations, a keep fraction of 0.4 and a mutation rate of 0.1. The minimization function was set as:

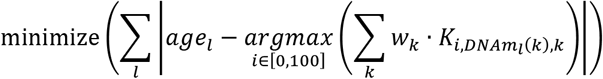

where *l* iterates for all the samples in the training set, *age*_*l*_ is the real age of the sample *l* and *DNAm*_*l*_(*k*) is the DNAm percentage at site *k* of the sample *l*. The genetic algorithm adjusts the weights of each CpG to minimize the sum of absolute differences between the predictions and the real ages, therefore finding the coefficients that minimize the total absolute error in the training set (Supplemental tables S2 and S3). After finding the optimal weights, the final predictions can be calculated as:

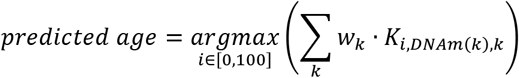

In a similar way, a probability vector can be calculated. Every vector has one entry for each year (0 to 100), which quantifies how probable it is to observe the measured methylation value assuming that a person is of the respective age. Since each CpG is associated with a weight coefficient, we have to add the individual weighed vectors for all the CpGs, which ends up in a single merged vector. The vectors can be subsequently normalized to interpret the weighted sum as a probability, i.e., to ensure that the sum of probabilities is equal to 1. This results in the following formula for weighted 2D KDE (WKDE model):

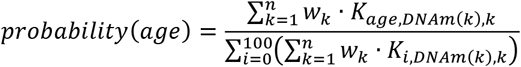

where *n* is the number of the CpGs that the clock uses, *w*_*k*_ is the weight of the CpG *k, DNAm*(*k*) is the measured DNAm percentage of the CpG *k* in the considered sample and *K*_*age*,*DNA*(*k*),*k*_ is the value in the row *DNAm*(*k*) and column *age* of the density map of the CpG *k*.

### Variation Score

The variation score provides an estimate for how homogeneous the age-predictions for individual CpGs are. Thus, we focus on the probability vectors themselves, without considering the weights. For this, we used the R function *approxfun* with the non-weighted probability function to obtain probability values for all ages from 0 to 100 in steps of 0.1 years. The function approxfun performs a linear interpolation between given data points:

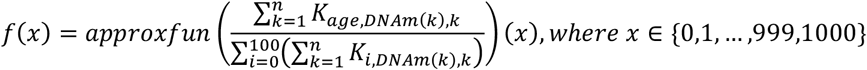

The call returns a vector with 1000 components, where the component x (which is between 0 and 1000) corresponds to the probability that the sample comes from an individual of age x/10. Subsequently, we were able to calculate mean, variance, and standard deviation of the probability function:

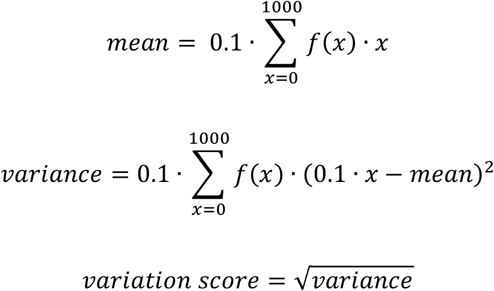

Finally, the Mann-Whitney-Wilcoxon Test was used in R with the function *wilcox*.*test* to verify if there were differences in delta age (predicted – chronological age) by sex.

### Epigenetic age predictions based on linear regression

To benchmark epigenetic age-predictions of our WKDE model with conventional approaches using the same subsets of CpGs (R^2^ > 0.6 or 0.7 in the training set, as indicated above), we established two alternative epigenetic clocks: 1) by multivariable linear regression using the *lm* function from stats package in R and the *predict* function to calculate age predictions for both training and validation sets (Supplemental Tables S2 and S3); and 2) by calculating the average of individual CpG age-predictions, whereby for each selected age-associated CpGs the slopes and intercepts between DNAm (beta values) and age were calculated with the lm and apply functions in R and the mean of the CpG-specific predictions was then used to estimate donor age (Supplemental Tables S2 and S3). Pearson squared correlation (R^2^) and median absolute error (MAE) between chronological and predicted age was calculated. Furthermore, we used the R package methylCIPHER (15) to compare the performance across the individual datasets used in training and validation sets with other commonly used epigenetic clocks: the Horvath clock (9), the Hannum clock (16), PhenoAge (17), retrained principal component PhenoAge (18), Horvath Skin & Blood epigenetic clock (19), Lin clock (20), Vidal-Bralo clock (21) and Han clock (22).

### Mortality association in LBC1921 and LBC1936

The Lothian Birth Cohorts of 1921 (LBC1921) and 1936 (LBC1936) are follow-up studies of the Scottish Mental Surveys of participants born in 1921 and 1936, respectively. The study was initially set up to study determinants of non-pathological cognitive aging (23, 24) and peripheral blood samples were analyzed by 450k Illumina BeadChips. Only samples from the first waves of the LBC1921 and LBC1936 were considered, in order to keep all the samples around the same chronological age (79.11 ± 0.59 years for LBC1921 and 69.56 ± 0.84 years for LBC1926) – thus, in these cohorts chronological age has little impact on the association with mortality analyses. All the raw data in the form of IDAT files were processed in R with the ENmix package. Quality control was performed, probes with detection p-values greater than 0.01 were filtered out, and probes and samples with more than 10% of missing values were excluded. ENmix_oob background correction (25), RELIC dye-bias correction (26) and RCP probe-type bias adjustment (27) were performed. To minimize the potential influence of fatal acute illnesses on the methylation measurements, deaths that occurred during the first two years of follow-up were excluded from the analysis. After the preprocessing filter for the first wave, a total of 374 samples from the LBC1921 and 721 from the LBC1936 remained.

Mortality status was ascertained from data linkage using dates of death (which were converted to age in days at death by the LBC research team) from the National Health Service Central Resister, provided by National Records of Scotland. Time of the events for the Cox models was defined as the interval between [*age in days at census wave1; age in days at death*] / 365.25 in case of a death or [*age in days at census wave1; age in days at last census*] / 365.25 in case of census. At time of last census, 367/374 (98.13%) and 285/721 (39.53%) had died. Delta age was calculated as predicted age (with the WKDE model) – chronological age. Cox proportional hazards regression models were performed (adjusted for age and sex), to assess the association between delta age and mortality and between variation score and mortality. The statistical software R was used to conduct all analyses, employing the ‘survival’ library for the Cox models.

## Results

### Two-dimensional kernel distribution provides probabilistic epigenetic age estimates

To determine epigenetic age based on KDE, we arbitrarily preselected CpGs with the highest linear correlation with chronological age in the training set (R^2^ > 0.7: 27 CpGs, or R^2^ > 0.6: 491 CpGs). For each preselected CpG the probability distribution of the age for a given DNAm level was determined by KDE and then combined into a joined probability estimation, where the age corresponding to the maximum probability was considered the predicted age of the sample. However, due to the heterogeneous distribution of donor ages, the predictions would be enriched at ages that are overrepresented in the training set (Figure 1A-E).

**Figure 1.**
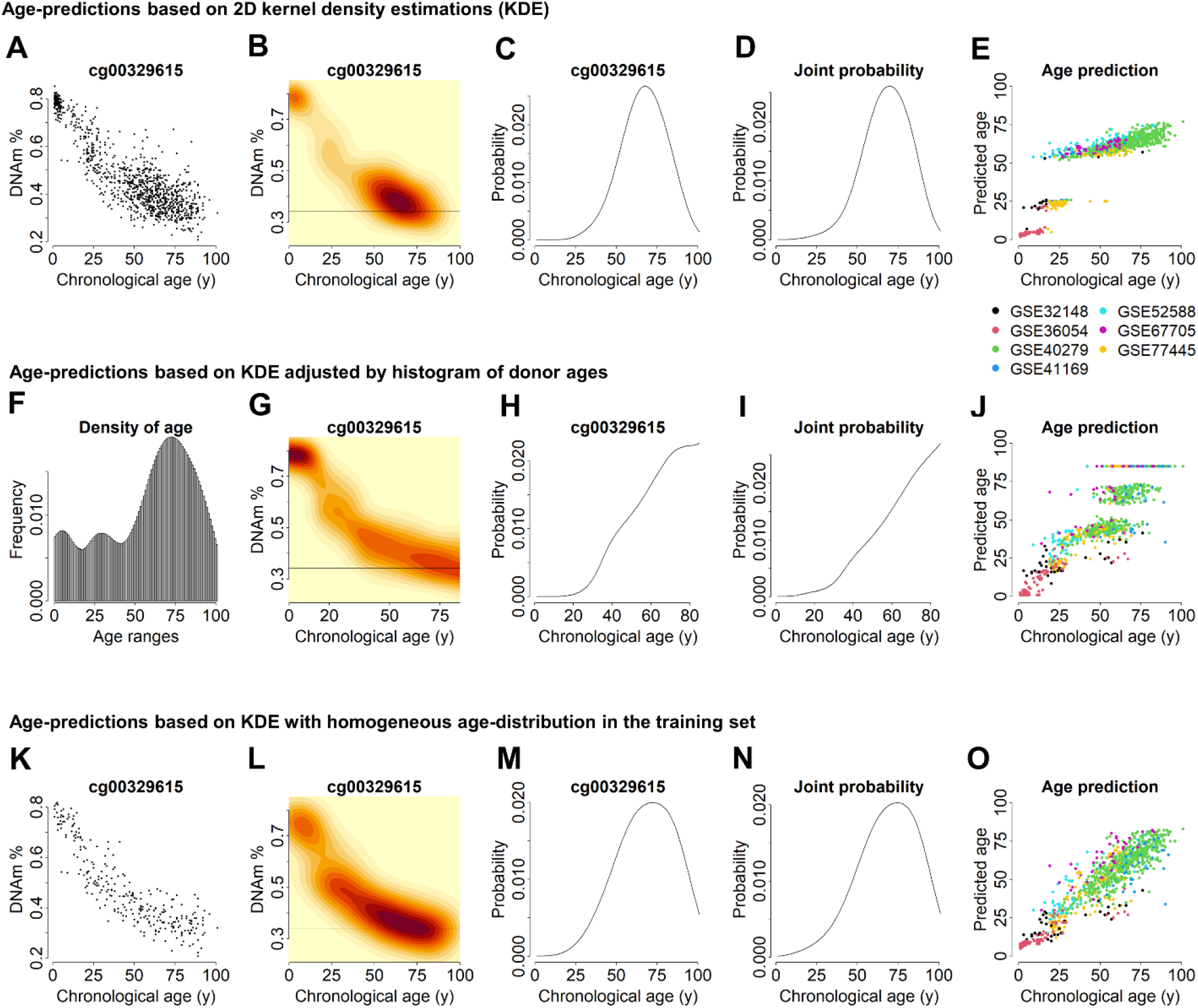
Construction of the probabilistic 2D kernel age predictor. **A)** The association of DNA methylation (DNAm) with chronological age is exemplarily depicted for the CpG sites cg00329615 for all the samples in the training set. **B)** 2D density kernel of the scatter plot depicted in A. The horizontal line exemplarily depicts the DNAm level of a given sample. **C)** Density distribution as estimated for the probability of age-predictions for the sample depicted in B. **D)** Joint probability for 27 age-associated CpGs for the same sample. **E)** Performance of the methodology shown A-D for all the samples in the training set (color code indicates different studies of the training set). **F)** Histogram of the ages in the training set. **G)** Normalization of 2D kernels by age (divided by the corresponding value in the histogram). **H)** Density distribution for the exemplary sample (line) in G. **I)** Joint probability 27 CpGs after normalization by age. **J)** Performance of the methodology shown F-I for all the samples in the training set. **K)** Alternatively, the training set was split in 5-year bins from 0 to 90 years (plus one bin of samples older than 90 years) and 15 samples from each bin were taken. The Scatter plot again depicts this selection for cg00329615. **L)** 2D density kernel of K. **M)** Density distribution for the sample in panel L. **N)** Joint probability for 27 CpGs of the same sample. **O)** Performance of the methodology shown K-N for all the samples in the training set.

Therefore, we normalized the KDE maps by the frequency of samples at each specific age. We excluded samples older than 85 years (N = 39), because their low frequency then resulted in high density in the 2D kernel maps. Adjustment of KDE by the histogram of donor ages provided more accurate predictions, but there were still clusters, indicating that the normalization now rather skewed for age-ranges with the highest density or with low number of samples (Figure F-J).

To further reduce the effects of age distribution in the training set we subsequently divided the samples into 18 bins of 5-year intervals, ranging from 0 to 90 years, plus one 19^th^ bin with the samples older than 90 years. From each bin, 15 samples were selected to construct the 2D density kernels (Supplemental Figure S1). This resulted in 2D density kernels with a more even distribution of densities and epigenetic age predictions that correlated with chronological age (Figure K-O). Thus, KDE can be used for a probabilistic approach for epigenetic age predictions.

### Improvement by weighted 2D kernel age-predictions

When we initially tested our probabilistic kernel models based on either 27 or 491 CpGs, correlation with chronological age in the training set was R^2^ = 0.86 or 0.79, respectively, with a median absolute error (MAE) of 5 or 6 years. However, in the independent validation set, the correlations and precisions of age-predictions were much lower (27 CpG model: R^2^ = 0.37 and MAE = 10 years, Figure 2A; 491 CpG model: R^2^ = 0.23 and MAE = 13 years, Supplemental Figure S2A), suggesting that there may be off-sets between different studies at individual CpGs, e.g. due to batch variation.

**Figure 2.**
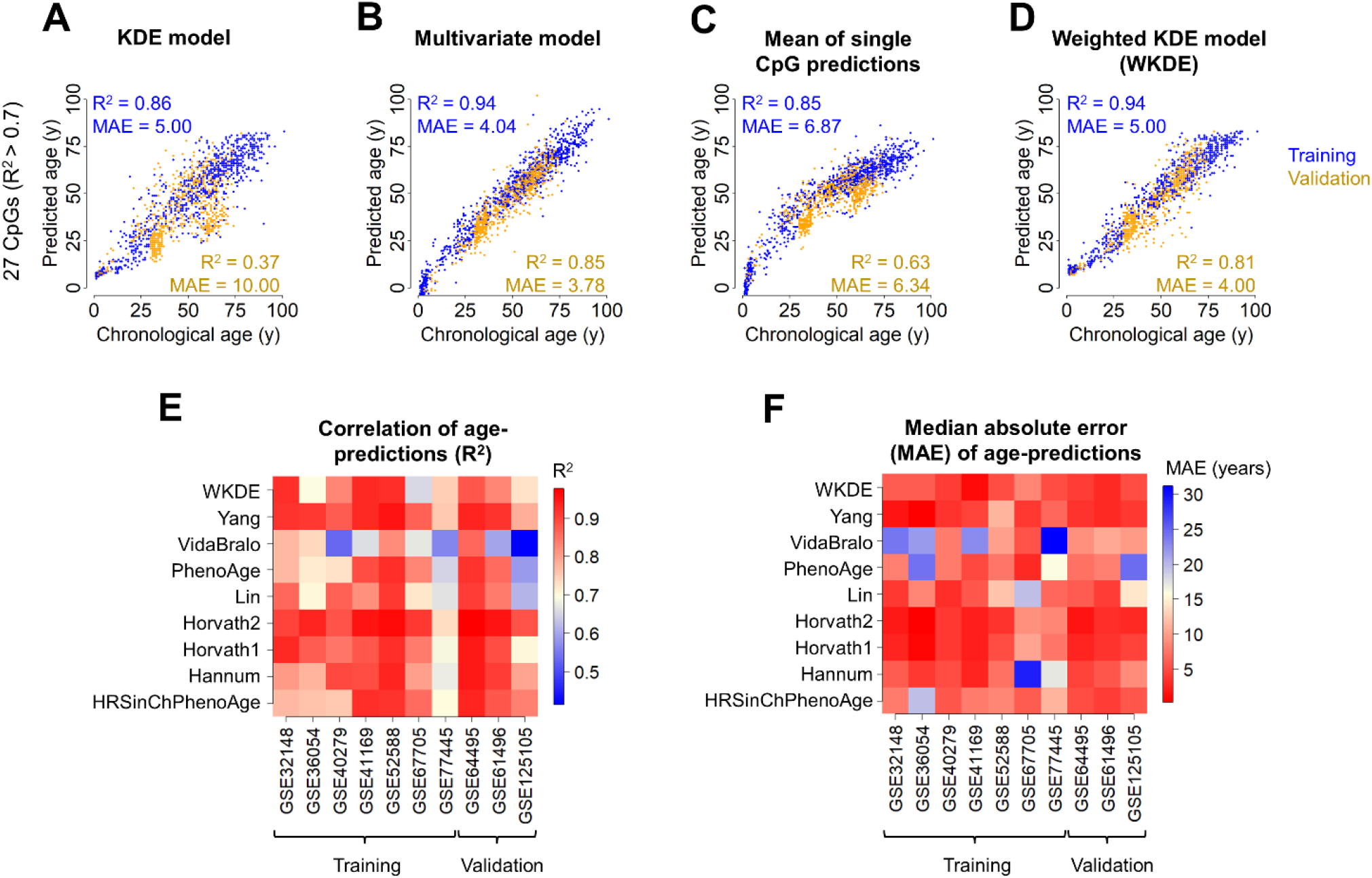
A weighted approach improves 2D kernel age predictions. **A)** 2D kernel age predication model was generated for 27 CpGs (R^2^ > 0.7 in the training set). The model was trained on a subset of samples from the training set with homogeneous age distribution. When we applied the model to the validation dataset, the predictions were not reliable. Pearson squared correlation R^2^ and median absolute error (MAE) are indicated. **B)** The same CpGs were used to generate multivariate models based on the entire training set. **C)** Alternatively, for each of the age-associated CpGs a line regression model was established to facilitate single CpG predictions. When we averaged these predictions, the performance was much lower than for the multivariable model. **D)** The 2D kernel age-prediction model was further optimized by optimized weights for individual CpGs, which were determined by genetic algorithm optimization. **E)** Heatmap of Pearson squared correlation (R^2^) between chronological and predicted age for WKDE and a set of widely used clocks, split in all datasets used for training and validation. **F)** Heatmap of MAE for WKDE and the same set of clocks as in **E**, split in all datasets used for training and validation.

For comparison, we used the same set of age associated CpGs to generate multivariate regression models. This approach provided higher correlation with chronological age in the training set (27 CpG model: R^2^ = 0.94, Figure 2B; 491 CpG model: R^2^ = 0.98, Supplemental Figure S2B). In fact, when creating multivariate models with the training set by varying the number of CpG sites, the highest correlation in the validation set was achieved with the top 125 CpGs (R^2^ = 0.92, Supplemental Figure S3A), while the highest precision in the validation set would have been achieved with the top 81 CpGs (MAE = 2.87 years, Supplemental Figure S3B), indicating that larger signatures are not always beneficial. In the independent validation set, the multivariable models with 27 or 491 CpGs revealed high correlation with age, too (both R^2^ = 0.85; Figure 2B, Supplemental Figure S2B). In contrast, when we just used the average of individual CpG predictions the age-estimates were much less precise (27 CpG model: training: R^2^ = 0.85, validation: R^2^ = 0.63, Figure 2C; 491 CpG model: training: R^2^ = 0.77, validation: R^2^ = 0.33, Supplemental Figure S2C). Furthermore, the ages of very young samples were overestimated due to exponential age-associated epigenetic changes in childhood (8). These findings supported the notion that using conventional regression approaches a multivariable weighted approach was clearly advantageous.

Consequently, we have also adjusted our probabilistic kernel models to better weight the impact of individual CpGs. To this end, we used genetic algorithm optimization to identify the best weights for each CpG according to the cumulative error curves. This weighted kernel density estimate model (WKDE) improved predictions, also in the validation set (Figure 2D, Supplemental Figure S2D) – particularly with the 27 CpG model the predictions (R^2^ = 0.94 and MAE = 5 years in the training set; R^2^ = 0.81 and MAE = 4 years in the validation set) were now in a similar range as observed for the multivariable model.

### Benchmarking with other commonly used epigenetic clocks

Subsequently, we compared the performance of the WKDE model with other widely used epigenetic clocks in the field. To better test the applicability to independent datasets, we have done this analysis for each study of the training and validation sets. The WKDE predictor revealed better or similar correlations to many other clocks, while particularly the Yang clock and Horvath Skin & Blood clock showed highest correlations (Figure 2E). Furthermore, the WKDE model provided small median absolute errors for all datasets – apart from Yang clock and both Horvath clocks, other clocks revealed high MAEs of more than 15 years for at least one dataset (Figure 2F). Thus, despite the small epigenetic signature of only 27 CpGs the WKDE model facilitated robust estimation of chronological age.

### Heterogeneity of age-associated DNAm within a given sample

Our probabilistic approach allows to calculate the probability of the age of a given sample for every year from 0 to 100 (Figure 3A). In addition, it provides an estimate of the disparity among the selected methylation sites, which is subsequently referred to as variation score. This measure reflects the heterogeneity in age-associated DNAm within our 27 CpG signature, rather than the probability that the estimates of chronological age are correct. However, for most of the samples in training set (99.22% of total samples; Figure 3B) and validation set (98.56% of total samples; Figure 3C) the chronological age falls into this range (predicted age ± variation score). The variation score shows small correlations with chronological age (training: R^2^ = 0.36, validation: R^2^ = 0.33), epigenetic age (training: R^2^ = 0.37, validation: R^2^ = 0.24), and delta age (training: R^2^ = 0.07, validation: R^2^ = 0.02) and was highest at an age-range from 30 to 75 years (Figures 3D-F). When we compared male and female samples, we did not observe any significant gender bias with regard to delta age (P < 0.37, Figure 3G), or variation scores (P < 0.92; Figure 3H).

**Figure 3.**
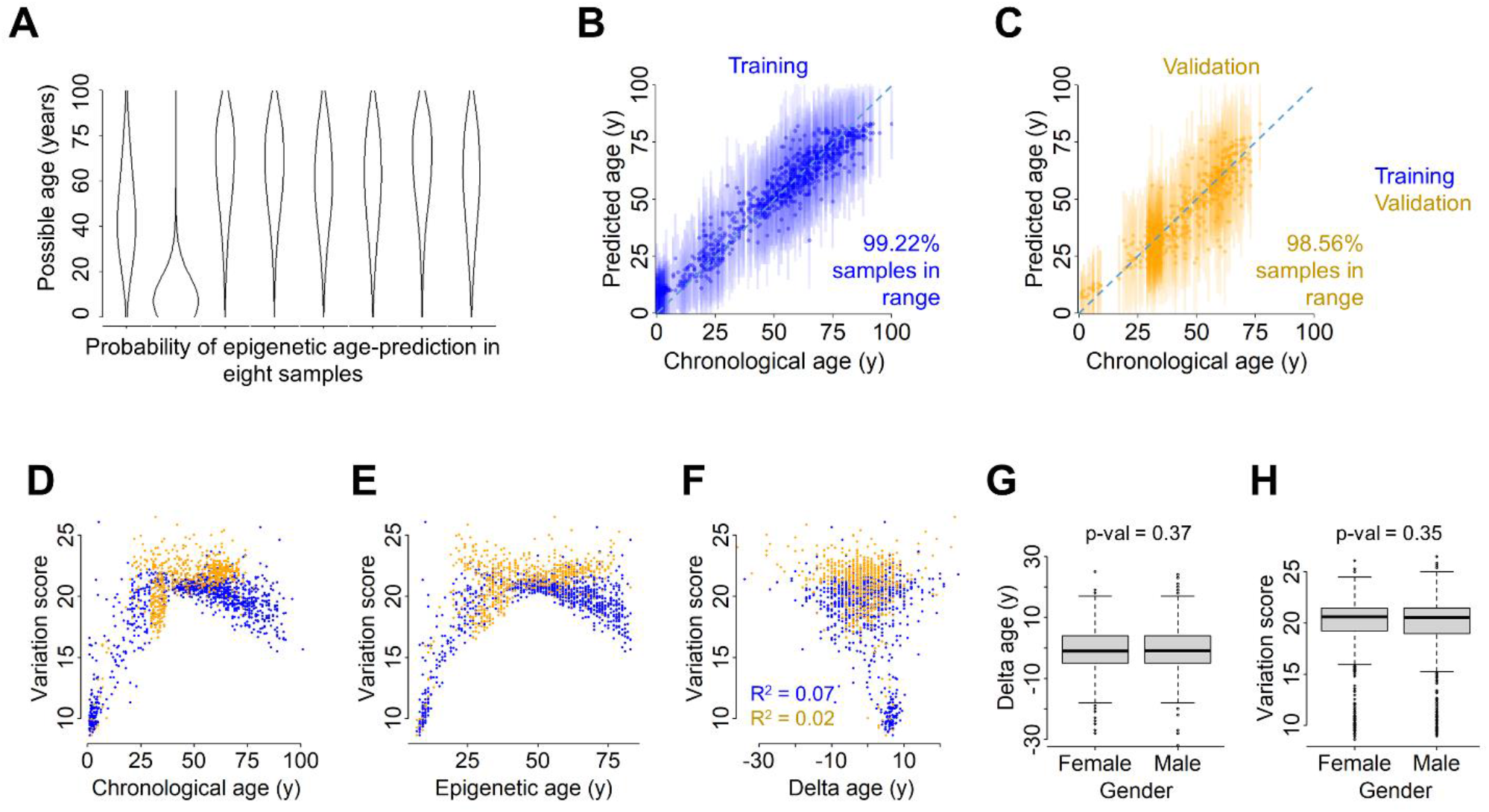
WKDE allows to measure heterogeneity in age-associated DNAm. **A)** Possible age (y-axis) vs. probability (x-axis) for eight random samples from the training set. The thickness of the plot represents how probable is that a sample belongs to that age. **B)** Chronological vs. predicted age in the training set measured with WKDE. Vertical lines depict predicted age ± variation score. **C)** Same as in panel B for validation set. **D)** Chronological age vs. variation score. **E)** Predicted age vs. variation score. **F)** Delta age vs. variation score. Variation score reflects almost no correlation with delta age measured with WKDE. **G)** Boxplot of delta age vs. gender. Delta age distributes evenly across males and females (Wilcoxon rank sum test P < 0.37). **H)** Boxplot of variation score vs. gender. Variation score distributes evenly across males and females (Wilcoxon rank sum test P < 0.35).

### Association of sample-intrinsic heterogeneity in epigenetic aging with all-cause mortality

It has been reported that all-cause mortality increases with accelerated epigenetic age in various clocks (5, 28). To estimate if this also applies for our WKDE model, we analyzed methylation from the first waves of the LBC1921 and LBC1936 with subsequent mortality risk. Cox proportional hazards regression models for delta age vs. survival, adjusting for age and sex, revealed no significant association between delta age and mortality in the LBC1921 (P < 0.978, Figure 4A) and in the LBC1936 (P < 0.415, Supplemental Figure S4A).

**Figure 4.**
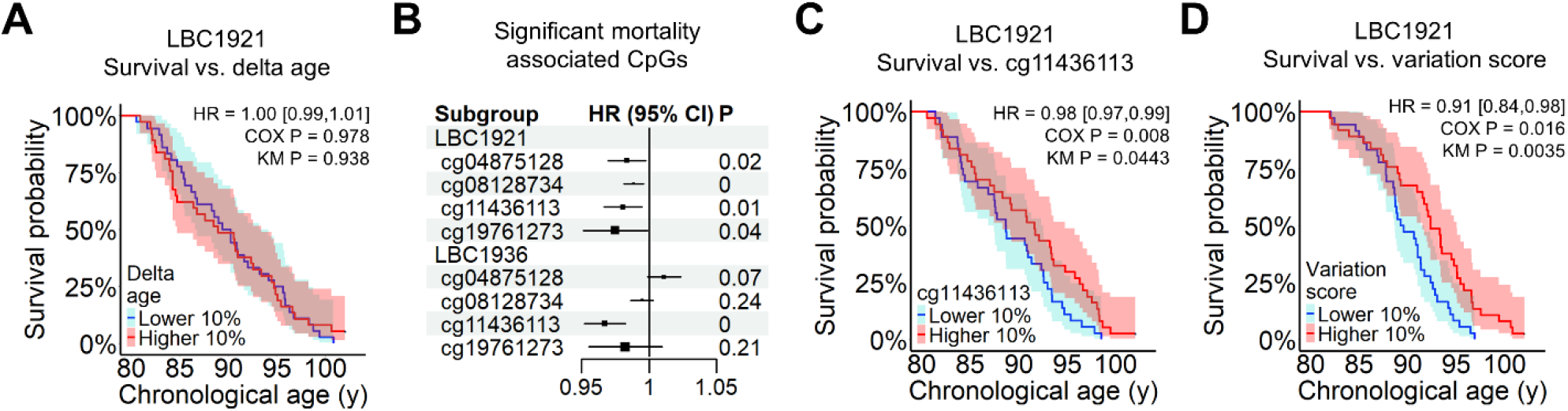
Variation score shows significant association with mortality. **A)** The association of epigenetic age-predictions with all-cause mortality was analyzed in the Lothian Birth Cohorts of 1921 (LBC1921). Kaplan-Meier survival curves are depicted for highest and lowest 10% delta ages. Cox regression model for all donors, adjusted for chronological age and gender, showed no significant effect of delta age in mortality risk (HR = 0.9998, 95% CI (0.988, 1.012), P <0.978). **B)** Individual Cox regressions for all the 27 CpGs in WKDE (adjusting for age and gender) revealed 4 significantly mortality-associated CpGs in the LBC1921. The cg11436113 was also significantly associated with mortality in the Lothian Birth Cohorts of 1936 (LBC1936). **C)** Kaplan-Meier survival curves for donors with highest and lowest 10% DNAm at cg11436113 in LBC1921. Cox regression model, adjusted for chronological age and gender, shows that increase of 1% in the DNAm of cg11436113 is associated with a 1,94% decrease in mortality risk (95% CI (0.9665, 0.9949), P < 0,0080). **D)** Kaplan-Meier survival curves for highest and lowest 10% variation score in LBC1921. Cox regression model, adjusted for chronological age and gender, shows that increase of 1 unit in the variation score is associated with a 9.2% decrease in mortality risk (95% CI (0.8387, 0.9872), P <0.0160).

Subsequently, we tested if any of the individual CpG sites in the 27 CpG or 491 CpG models were associated with all-cause mortality, after adjustment for age and gender. Within the 491 CpGs we found 25 mortality associated sites that overlap between LBC1921 and LBC1936 (Supplemental Figure S4B and Supplemental Tables S4 and S5), with almost all of them being hypomethylated with aging. Out of the 27 CpG sites, four were significantly associated with mortality in the LBC1921 and one in the LBC1936 (Figure 4B). Particularly cg11436113, which is located in an intergenic region, was highly associated with mortality in both cohorts (LBC1921: P < 0.008, Figure 4C; LBC1936: P < 0.00003, Supplemental Figure S4C).

When testing for associations between variation score and mortality in the LBC1921 cohort, we observed that an increase of 1 unit in the variation score is associated with a 9.2% decrease in mortality risk (95% CI (0.8387, 0.9872), P < 0.0160, Figure 4D), after adjusting for gender and chronological age. The results remained even significant when we adjusted for gender, chronological age, and delta age (P = 0.0159), indicating that the variation score might be an independent measure for biological aging. However, a significant association of the variation score with all-cause mortality was not observed in the LBC1936 cohort (HR = 0.9894, 95% CI (0.8870, 1.1037), P < 0.849, Supplemental Figure S4D), which might partly be attributed to a lower number of deaths in this cohort.

## Discussion

In this study, we demonstrate that 2D kernel density estimation can be used for robust epigenetic age-predictions. Age-associated DNAm does not necessarily follow a strictly linear or logarithmic pattern (29). It is therefore advantageous that WKDE models are taking the entire data distribution into account. In fact, we did not observe over-estimation of epigenetic age in children with WKDE, whereas a logarithmic transformation is usually required for conventional multivariable clocks on pediatric samples (8, 9).

Nonetheless, WKDE has some limitations. Predictions are only possible within the age-range that is covered by the training set and the estimated kernels – in our case, we can therefore not reliably predict ages above 100 years. The imbalance of different donor-ages in the training set was a major problem, which we could neither entirely corrected by normalization with the age-distribution, nor by a conditional density-based approach for the creation of the 2D density kernels (14). We have therefore randomly reduced the training set for a homogeneous age-distribution – with the downside that not the entire training set was considered. In the future, alternative approaches based on resampling or machine learning might be further exploited to solve this limitation (30, 31). Furthermore, our WKDE model was optimized with a genetic algorithm, which has shown several advantages over other non-linear optimization techniques, such as high efficiency, parallelization and no need of derivative information (32). However, other tuning parameters for the genetic algorithm or even non-linear optimization algorithms can be tested in subsequent studies.

Larger epigenetic signatures do not always increase the precision. When we only considered the top 27 CpGs to build the WKDE model, we observed a higher correlation with chronological age and lower median absolute error than using a less stringent pre-selection of 491 CpGs. Theoretically, automatic CpG selection approaches, such as ElasticNet (33), could be used to further optimize the selection of CpGs for WKDE – in analogy to the ElasticNet based conventional epigenetic clocks (9, 16) – but it would require much more computational power to run it with WKDE as compared to the linear approaches. Our model was specifically trained for human blood samples, while similar models might also be trained for other cell types or tissues in the future. On the other hand, the 2D kernel density approach might also be applied for other epigenetic biomarkers, such as deconvolution of leukocytes in blood (34), or of cell types within a tissue (35, 36).

Probabilistic approaches for epigenetic clocks have been described before. For example, we have previously described probabilistic epigenetic age predictions for individual sequencing reads, using the binary sequel of methylated and non-methylated CpGs in barcoded bisulfite amplicon sequencing data (22). This method has been further developed for genome wide single cell DNAm datasets (37). In contrast, WKDE is not applicable to shallow sequencing data, but it provides density estimates for each CpG, a joint probability range for the sample and a variability score associated to it.

Although association between accelerated age and mortality has been shown before (5, 28), we did not observe this with WKDE in the LBC1921, which might be due to the sample selection and the nature of the clock itself. We have intentionally only chosen samples from the initial waves of the LBC1921 and LBC1936 to isolate the effects of the variables under examination on mortality from the confounding factor of aging (which differs from the previously mentioned studies). It has to be noted, that our WKDE model was trained to correlate as closely as possible with chronological age and it is therefore conceivable that mortality-associated CpGs sites – such as cg11436113 – were considered with a negative weight for age-predictions. Furthermore, WKDE employs a method of prediction that focuses on determining the age that maximizes the probability of a joint probability function for the age, rather than relying on a singular value. Hence, the biological age might still fall into the probability range of the age.

We have previously identified individual CpG sites from our 99-CpG clock and other epigenetic clocks that showed correlations with survival in both the LBC1921 and LBC1936 (20) – and most of these were hypomethylated with age (38). Here, we showed an overlap of 25 methylation sites that are significantly associated with mortality in both cohorts, and again most of these CpGs become hypomethylated with age. Out of these 25 CpGs, 7 are located in genes *BST2, FKBP5, PRDX5, NWD1, FAM38A* and *NOD2*, which have previously shown high association between DNAm and aging (39-42), and we have already identified cg16363586 (located in gene *BST2*) as life expectancy associated site (20). In addition, the CpG site that was identified as mortality-associated in both cohorts with the 27 CpG signature, cg11436113, has been reported as smoking and cancer related (43, 44).

While previous studies have discussed the link between age-related DNA methylation (DNAm) and mortality (45, 46) the connection between age-related DNAm heterogeneity and all-cause mortality was so far not addressed. Our results suggest that heterogeneity among age-associated methylation sites in the genome, measured as variability score, can provide additional insight into biological age. Nevertheless, further validation is necessary to examine the potential application of the variability score to estimate life-expectancy and to explore possible association with specific diseases.

## Conclusions

Our study introduces weighted 2D-kernel density estimation (WKDE) to perform accurate epigenetic age predictions. Furthermore, we describe a variation score that may act as an additional parameter for evaluating biological age.

## Declarations

### Ethics approval and consent to participate

LBC1936 ethical approval was obtained from the Multicentre Research Ethics Committee for Scotland (baseline, MREC/01/0/56), the Lothian Research Ethics Committee (age 70, LREC/2003/2/29), and the Scotland A Research Ethics Committee (ages 73, 76, 79, 82, 07/MRE00/58). LBC1921 Ethical approval was provided by the Lothian Research Ethics Committee for test waves 1–3 at ages 79, 83 and 87 (LREC/1998/4/183, LREC/2003/7/23, 1702/98/4/183) and the Scotland A Research Ethics Committee for test wave 4 at age 90 (10/MRE00/87, 10/MRE00/87). All participants provided written informed consent.

## Consent for publication

Not applicable

## Availability of data and materials

The DNAm and gene expression datasets analyzed in this study are available in NCBI’s Gene Expression Omnibus repository (https://www.ncbi.nlm.nih.gov/geo/) under the accession numbers as indicated in the text. Information on data availability for the Lothian Birth Cohorts (along with a template data request form and data dictionaries) can be found at the study website: https://lothian-birth-cohorts.ed.ac.uk/data-access-collaboration.

## Authors’ contributions

J-F.P-C. implemented, adapted and/or created the prediction models, performed the analysis and created the figures; T.S. and I.G.C. supervised and helped in the development of the algorithm; R.E.M., J.C. and S.R.C. supported analysis of the LBC cohort; W.W. and J-F.P-C. wrote the manuscript. All authors contributed and approved the final version.

## Supplemental Data

**Additional file 1: All supplemental figures and tables.**

