## Supplemental Figures and Tables for "Weighted 2D-kernel density estimations provide a new probabilistic measure for epigenetic age"

**Supplemental file 1**

### Weighted 2D-kernel density estimations provide a new probabilistic measure for epigenetic age

Juan-Felipe Perez-Correa, Thomas Stiehl, Riccardo E. Marioni, Janie Corley, Simon R. Cox, Ivan Costa, Wolfgang Wagner

#### Index

#### Supplemental Figures

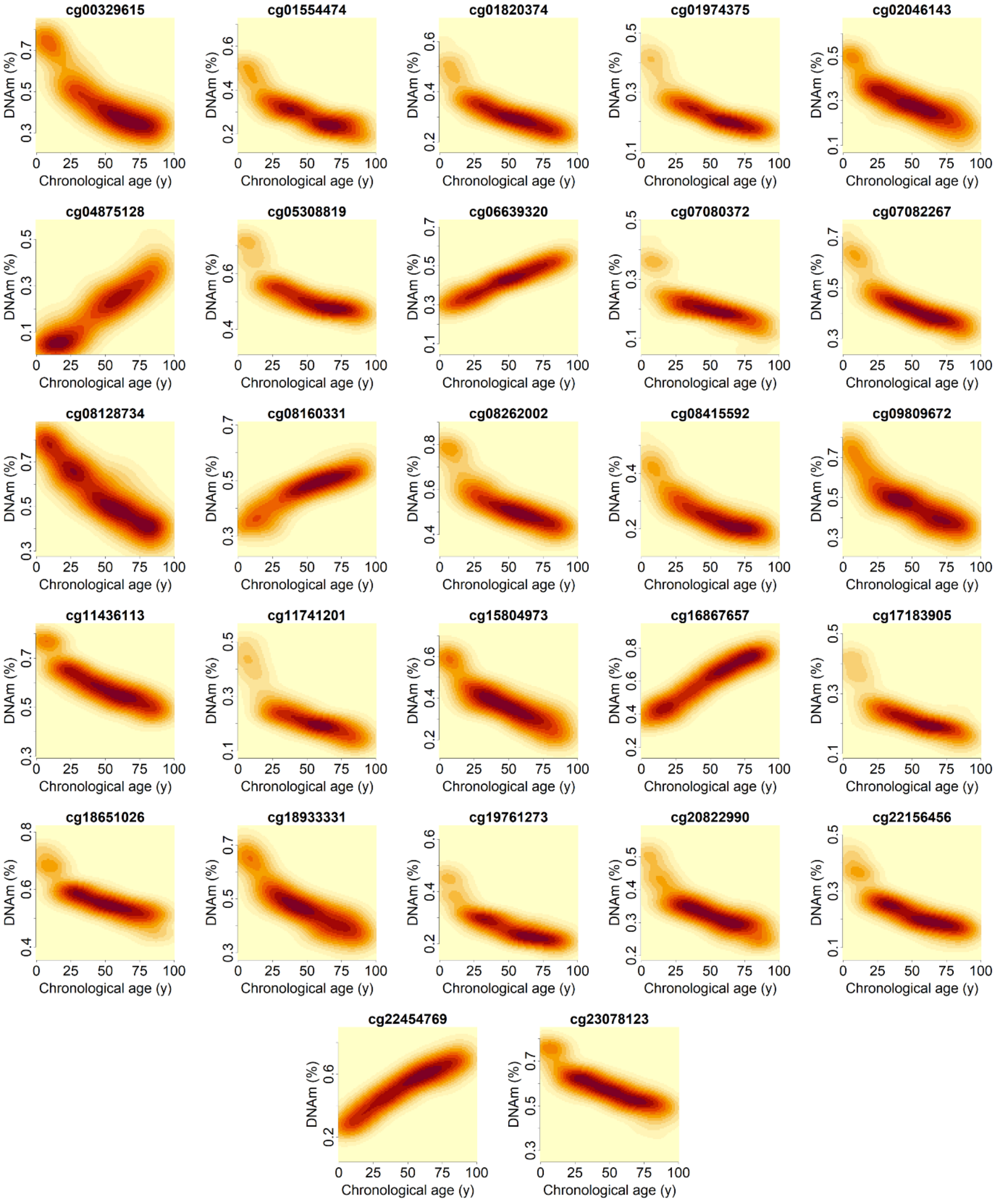

**Supplemental Figure S1. Kernel density maps of the 27 age-associated CpGs.**
For the 27 age-associated CpGs the 2D density kernel estimations are depicted for a homogeneous age-distribution in the training set.

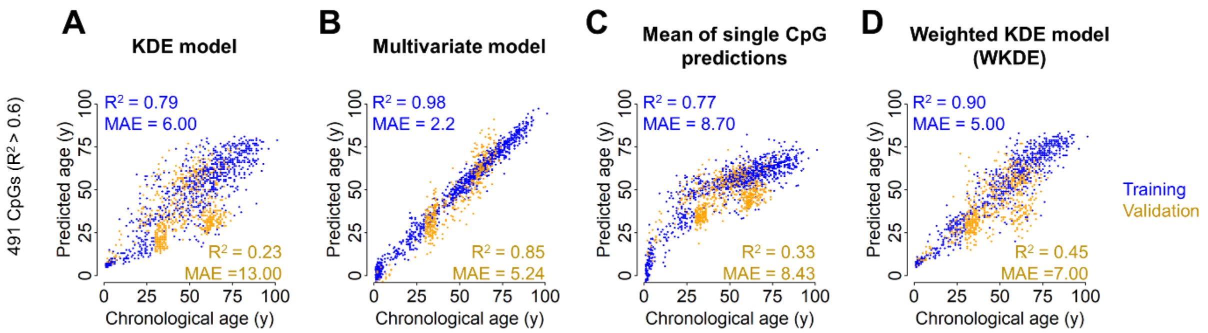

**Supplemental Figure S2. Weighted approach with 491 CpG model.**

**A)** An alternative 2D kernel age predication model was generated for 491 CpGs ($R^2 > 0.6$ in the training set). The model was trained on a subset of samples from the training set with homogeneous age distribution. Pearson correlation $R^2$ and mean absolute error (MAE) are indicated either for all data of the training sets (blue; even though most of the samples were not selected for the homogeneous age presentation of the training data), and the independent validation sets (yellow). **B)** The same CpGs were used to generate multivariate models based on the entire training set. **C)** Alternatively, for each of the 491 age-associated CpGs a line regression model was established to facilitate single CpG predictions. The averaged of these predictions revealed a much lower precision of age predictions than for the multivariable model. **D)** The 491 CpG 2D kernel age-prediction model was further optimized by optimized weights for individual CpGs that were determined by genetic algorithm optimization. However, the performance in the validation set is lower than the one with the 27 CpG model.

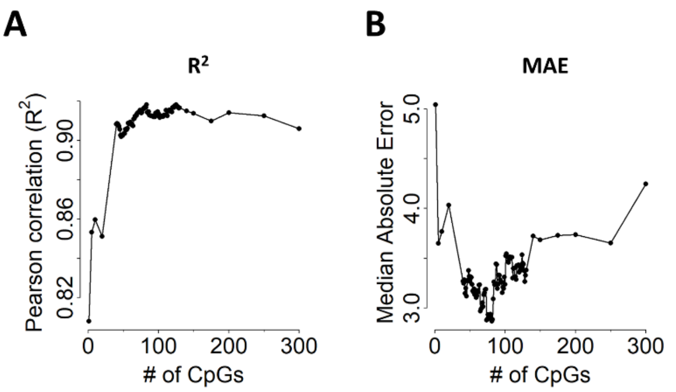

**Supplemental Figure S3. Multivariate linear models with different numbers of CpGs.**
**A)** To test how the performance of multivariate linear models depends on the number of age-associated CpGs we trained such models with up to the top 300 age-associated CpGs in the training set. The highest Pearson correlation with chronological age was achieved with the model using 125 CpGs ($R^2$ = 0.92). **B)** Median absolute errors of the models mentioned in A. The smallest error was observed using the top 81 CpGs (MAE = 2.87 years).

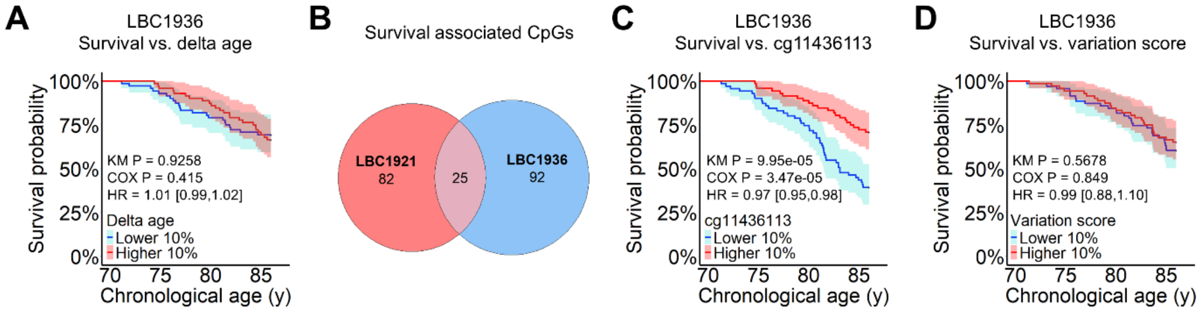

**Supplemental Figure S4. Mortality analysis in LBC1936.**
**A)** The association of epigenetic age-predictions of the 27 WKDE model with all-cause mortality was also tested in the Lothian Birth Cohorts of 1936 (LBC1936). Kaplan-Meier survival curves for donors with highest and lowest 10% delta ages are depicted. Cox regression model for all donors, adjusted for chronological age and gender, showed no significant effect of delta age in mortality risk (HR = 1.005, 95% CI (0.9923, 1.0183), P < 0.415). **B)** Individual Cox regressions for all the 491 age-associated CpGs (adjusting for age and gender) revealed high overlap in mortality associated CpG sites in both cohorts. **C)** Kaplan-Meier survival curves for highest and lowest 10% methylation values of cg11436113 in LBC1936. Cox regression model, adjusted for chronological age and gender, shows that increase of 1% in the DNAm of cg11436113 is associated with a 3.30% decrease in mortality risk (95% CI (0.9520, 0.9830), P < 3.47E-5). **D)** Kaplan-Meier survival curves for highest and lowest 10% variation score in LBC1936. Cox regression model, adjusted for chronological age and gender, showed no significant effect of the variation score in mortality risk in the LBC1936 (HR = 0.9894, 95% CI (0.8870, 1.1037), P <0.849).

### Supplemental Tables

**Supplemental Table S1. Illumina BeadChip profiles used for training and validation sets.**

| GEO Accession | Number of samples | Number of healthy or control | Age range healthy or control | Gender distribution (male%) | Tissue of samples | Set |
|---|---|---|---|---|---|---|
| GSE36054 | 192 | 134 | 1-17 y | 58.96% | Leukocyte | Training |
| GSE32148 | 48 | 19 | 3-76 y | 36.84% | Periph. Blood | Training |
| GSE41169 | 95 | 33 | 18-65 y | 63.64% | Whole blood | Training |
| GSE77445 | 85 | 85 | 18-69 y | 50.59% | Whole blood | Training |
| GSE52588 | 87 | 58 | 9-83 y | 28.74% | Whole blood | Training |
| GSE67705 | 284 | 44 | 27-66 y | 65.49% | Whole blood | Training |
| GSE40279 | 656 | 656 | 19-101 y | 48.48% | Whole blood | Training |
| **Total Training** | **1447** | **1029** | **1-101 y** | **49.93%** | | |
| GSE64495 | 113 | 106 | 2-73 y | 32.74% | Whole blood | Validation |
| GSE61496 | 312 | 312 | 30-74 y | 52.56% | Whole blood | Validation |
| GSE125105 | 699 | 210 | 19-79 y | 44.64% | Whole blood | Validation |
| **Total Validation** | **1124** | **628** | **2-79 y** | **46.57%** | | |

**Supplemental Table S2. List of 27 age-associated CpGs and variables for different models.**

| CpG | Coefficients of multivariate models | Slope of individual linear regression | Intercept of individual linear regression | Weights of WKDE | CHR | LOC | Strand | UCSC Ref. Gene Name |
|---|---|---|---|---|---|---|---|---|
| (Intercept) | 34,77 | - | - | - | | | | |
| cg00329615 | -11,01 | -145,07 | 118,47 | 2,21 | 3 | 118706648 | - | *CASZ1* |
| cg01554474 | -13,42 | -214,62 | 117,38 | 1,73 | 1 | 155107599 | + | *DGKI* |
| cg01820374 | 8,61 | -223,49 | 123,03 | 0,72 | 12 | 6882083 | - | |
| cg01974375 | 21,81 | -250,97 | 112,92 | -0,82 | 1 | 151298954 | + | |
| cg02046143 | -5,03 | -193,35 | 106,50 | 6,36 | 11 | 133797911 | + | *SMAP1* |
| cg04875128 | 31,77 | 168,76 | 10,48 | 6,95 | 15 | 31775895 | - | *HDAC4* |
| cg05308819 | -38,35 | -241,64 | 178,70 | 1,96 | 1 | 155959156 | - | *HINT2* |
| cg06639320 | 86,65 | 252,91 | -59,85 | 6,41 | 2 | 106015739 | - | |
| cg07080372 | -47,81 | -267,00 | 107,14 | 3,38 | 11 | 796607 | - | *IGSF11* |
| cg07082267 | 13,11 | -209,06 | 142,41 | 2,03 | 16 | 85429035 | - | *CASS4* |
| cg08128734 | -13,57 | -156,31 | 138,19 | 2,34 | 1 | 206685423 | - | *PTGDS* |
| cg08160331 | 4,02 | 274,45 | -76,25 | 2,45 | 11 | 75140865 | + | |
| cg08262002 | 9,84 | -184,95 | 151,48 | 1,03 | 4 | 16575323 | + | *ZEB2* |
| cg08415592 | -18,56 | -228,55 | 113,12 | 1,85 | 22 | 36648973 | - | *ZEB2* |
| cg09809672 | -13,55 | -163,69 | 131,16 | 3,69 | 1 | 236557682 | - | *TCEA2* |
| cg11436113 | -13,00 | -234,05 | 189,57 | 3,47 | 20 | 19191145 | - | |
| cg11741201 | 3,38 | -210,59 | 102,49 | -0,65 | 11 | 35638398 | - | *TJP2* |
| cg15804973 | -3,36 | -177,55 | 118,91 | 1,29 | 6 | 137114513 | + | *MMEL1* |
| cg16867657 | 55,03 | 173,46 | -55,16 | 7,63 | 6 | 11044877 | + | |
| cg17183905 | -18,78 | -242,45 | 107,33 | 1,26 | 12 | 110253842 | - | |
| cg18651026 | 14,14 | -270,92 | 203,20 | 0,28 | 6 | 33140660 | - | *DDO* |
| cg18933331 | -17,34 | -210,85 | 151,07 | 1,42 | 1 | 110186418 | - | *NANOS3* |
| cg19761273 | 20,50 | -261,03 | 123,42 | -3,47 | 17 | 80232096 | + | *CD300LB* |
| cg20822990 | 9,64 | -270,87 | 143,67 | -2,01 | 1 | 17338766 | - | |
| cg22156456 | 26,88 | -247,48 | 108,72 | -0,42 | 17 | 39844239 | - | *SNX20* |
| cg22454769 | -24,50 | 149,85 | -29,55 | 3,79 | 2 | 106015767 | - | *MIR770;MEG3* |
| cg23078123 | -22,03 | -232,69 | 187,66 | -0,05 | 1 | 68577796 | + | *EHD2* |

For each of the 27 CpGs, we include the coefficients for the multivariate model, the slopes and intercepts for the average of n-independent linear regressions, and the weights for the WKDE model.

**Supplemental Table S3. List of 491 age-associated CpGs and variables for different models.**

| CpG | Coefficients of multivariate model | Slopes of individual regression | Intercepts of indiv. reg, | Weight of WKDE model | CHR | LOC | Strand | UCSC Ref. Gene Name |
|---|---|---|---|---|---|---|---|---|
| (Intercept) | 21,88 | | | | | | | |
| cg00003345 | 0,02 | -294,32 | 231,35 | 0,70 | 1 | 10816499 | - | *CASZ1* |
| cg00029246 | 14,34 | -149,13 | 121,16 | 1,07 | 7 | 137223417 | + | *DGKI* |
| cg00101260 | 6,52 | -247,53 | 122,71 | 1,20 | 17 | 53426657 | - | |
| cg00103778 | 3,83 | -217,65 | 112,68 | 1,59 | 20 | 30196041 | + | |
| cg00129827 | 0,76 | -237,60 | 106,88 | 0,26 | 6 | 71394664 | + | *SMAP1* |
| cg00144180 | 32,77 | 218,95 | -134,16 | 2,16 | 2 | 240294362 | - | *HDAC4* |
| cg00193668 | 3,51 | -255,58 | 188,14 | -0,87 | 9 | 35814283 | + | *HINT2* |
| cg00232105 | -1,66 | -274,93 | 211,60 | 5,79 | 14 | 100752338 | + | |
| cg00329615 | 1,51 | -145,07 | 118,47 | 2,39 | 3 | 118706648 | - | *IGSF11* |
| cg00387658 | -10,49 | -236,71 | 182,05 | -1,25 | 20 | 54986793 | + | *CASS4* |
| cg00563932 | -6,61 | -218,23 | 158,22 | 5,68 | 9 | 139871049 | + | *PTGDS* |
| cg00570618 | 8,16 | -211,86 | 84,53 | 0,04 | 8 | 81143035 | + | |
| cg00573770 | -0,18 | -133,82 | 104,46 | 2,31 | 2 | 145278485 | + | *ZEB2* |
| cg00602811 | -7,13 | -141,70 | 113,11 | 3,29 | 2 | 145278564 | + | *ZEB2* |
| cg00636737 | 0,06 | -287,28 | 259,41 | -2,85 | 20 | 62695569 | - | *TCEA2* |
| cg00655552 | 7,28 | -209,87 | 73,30 | -3,56 | 4 | 182862370 | - | |
| cg00695112 | 16,13 | -269,01 | 222,26 | 0,52 | 9 | 71862747 | - | *TJP2* |
| cg00695391 | 4,73 | -177,87 | 118,04 | -2,20 | 1 | 2525548 | - | *MMEL1* |
| cg00753885 | -8,64 | -352,74 | 310,14 | 3,56 | 12 | 57401797 | - | |
| cg00792107 | -7,15 | -223,29 | 119,29 | 1,35 | 12 | 133181600 | - | |
| cg00804078 | 3,24 | -124,53 | 96,74 | -1,73 | 6 | 110736941 | + | *DDO* |
| cg00863306 | -1,53 | -301,57 | 96,15 | -6,56 | 19 | 13991470 | + | *NANOS3* |
| cg00873351 | 11,21 | -193,91 | 125,32 | -0,24 | 17 | 72527813 | + | *CD300LB* |
| cg00876267 | 12,39 | -256,34 | 210,19 | 3,69 | 9 | 139588516 | + | |
| cg00921350 | 1,33 | -253,07 | 194,05 | -3,64 | 16 | 50701499 | - | *SNX20* |
| cg01022345 | -19,99 | -292,83 | 232,39 | -7,15 | 14 | 101317819 | + | *MIR770;MEG3* |
| cg01055871 | 1,74 | -291,65 | 120,98 | 0,41 | 19 | 48216390 | + | *EHD2* |
| cg01102833 | -3,60 | -253,36 | 115,79 | -2,00 | 4 | 176986621 | - | *WDR17* |
| cg01102854 | -16,03 | -234,09 | 174,69 | 1,36 | 11 | 63605717 | + | *MARK2* |
| cg01214061 | -5,06 | -295,45 | 79,75 | 0,78 | 4 | 174290520 | - | |
| cg01234420 | 9,90 | -154,57 | 124,71 | 1,98 | 22 | 46453808 | + | *LOC150381* |
| cg01243823 | 7,40 | -161,72 | 103,98 | -1,00 | 16 | 50732212 | - | *NOD2* |
| cg01282174 | 5,98 | -130,52 | 84,37 | -1,20 | 11 | 119630144 | + | |
| cg01314044 | 2,75 | -206,84 | 110,33 | 2,60 | 11 | 94246036 | + | |
| cg01447660 | -13,11 | -236,84 | 120,25 | -1,74 | 2 | 69152678 | - | |
| cg01502244 | -26,15 | -361,72 | 317,41 | 5,42 | 17 | 78188904 | + | *SGSH* |
| cg01506917 | 3,23 | -181,39 | 130,47 | -2,34 | 6 | 42417086 | - | *TRERF1* |
| cg01511567 | 1,83 | -267,54 | 109,63 | 4,67 | 11 | 57103631 | + | *SSRP1* |
| cg01514353 | 12,84 | -227,11 | 118,43 | 5,02 | 22 | 46259643 | - | |
| cg01528542 | -14,06 | -218,48 | 185,24 | 5,69 | 12 | 81468232 | + | |
| cg01538166 | 25,68 | -183,54 | 99,01 | 1,42 | 17 | 17743987 | + | |
| cg01542019 | -6,78 | -162,25 | 137,60 | 0,57 | 19 | 14673053 | - | *TECR* |
| cg01554474 | -14,00 | -214,62 | 117,38 | 2,25 | 1 | 155107599 | + | *RAG1AP1* |

| | | | | | | | | |
|---|---|---|---|---|---|---|---|---|
| cg01594949 | 5,88 | -239,86 | 190,51 | -2,28 | 9 | 140197890 | + | *NRARP* |
| cg01649334 | -10,00 | -236,58 | 97,39 | 3,38 | 3 | 101547296 | + | *NFKBIZ* |
| cg01676322 | 21,15 | -443,73 | 117,39 | -0,74 | 17 | 43213429 | - | *ACBD4* |
| cg01719405 | -12,52 | -169,81 | 120,33 | 4,37 | 14 | 62401258 | + | |
| cg01812894 | -0,92 | -203,72 | 167,12 | 8,88 | 9 | 75568506 | - | *ALDH1A1* |
| cg01820374 | 1,36 | -223,49 | 123,03 | 0,58 | 12 | 6882083 | - | *LAG3* |
| cg01881062 | -5,41 | -202,29 | 119,73 | -0,61 | 1 | 6660403 | + | *KLHL21* |
| cg01910639 | -24,17 | -254,33 | 207,95 | 1,92 | 1 | 153507779 | - | *S100A6* |
| cg01955153 | 17,37 | -271,17 | 101,76 | -6,88 | 16 | 50769852 | + | |
| cg01974375 | -8,76 | -250,97 | 112,92 | 0,84 | 1 | 151298954 | + | *PI4KB* |
| cg01981760 | -2,86 | -423,95 | 117,46 | -1,72 | 16 | 53737576 | + | *FTO;RPGRIP1L* |
| cg02010481 | 3,25 | -158,13 | 85,54 | -4,57 | 7 | 28218524 | - | *JAZF1* |
| cg02030542 | -8,10 | -223,01 | 174,64 | 6,60 | 16 | 70722454 | + | *VAC14* |
| cg02046143 | 4,31 | -193,35 | 106,50 | 2,86 | 11 | 133797911 | + | *IGSF9B* |
| cg02151301 | 7,17 | -276,14 | 146,28 | 8,57 | 20 | 30101785 | - | *HM13* |
| cg02286081 | -2,28 | -180,56 | 86,08 | 0,41 | 6 | 33043841 | - | *HLA-DPB1* |
| cg02361903 | -7,32 | -214,84 | 145,95 | -0,40 | 1 | 110452616 | - | *CSF1* |
| cg02395812 | 5,16 | -332,13 | 159,08 | -2,60 | 14 | 105955879 | + | *C14orf80* |
| cg02402091 | -5,10 | -218,54 | 150,84 | 0,52 | 10 | 114135092 | + | *ACSL5* |
| cg02610723 | -37,22 | -330,23 | 97,50 | 7,64 | 16 | 88850534 | - | *FAM38A* |
| cg02797271 | 18,65 | -311,15 | 95,41 | 1,56 | 14 | 105532159 | + | *GPR132* |
| cg03039990 | 15,05 | -193,86 | 126,97 | 0,80 | 14 | 101317622 | - | *MIR770;MEG3* |
| cg03043157 | -3,54 | -136,40 | 94,00 | 1,93 | 6 | 97010903 | + | *FHL5* |
| cg03055693 | 5,14 | -282,61 | 235,64 | -5,86 | 20 | 60910388 | - | *LAMA5* |
| cg03172991 | 36,91 | -325,71 | 216,59 | 1,72 | 19 | 13105728 | + | *NFIX* |
| cg03206537 | 6,34 | -199,68 | 100,61 | 4,34 | 20 | 44521739 | + | *CTSA* |
| cg03211864 | 8,94 | -271,93 | 127,34 | -0,74 | 10 | 124060833 | + | *BTBD16* |
| cg03359362 | 4,68 | -338,44 | 84,27 | -1,04 | 19 | 47289611 | - | *SLC1A5* |
| cg03431918 | 4,33 | -236,94 | 94,41 | -3,99 | 17 | 77716367 | - | |
| cg03443986 | -15,08 | -308,88 | 94,33 | 1,59 | 2 | 65100572 | + | |
| cg03474926 | 32,86 | -340,34 | 181,38 | 1,06 | 9 | 136023407 | - | *RALGDS* |
| cg03530962 | -3,72 | -239,81 | 202,14 | 2,16 | 3 | 58475833 | - | |
| cg03551243 | 2,08 | -389,39 | 92,04 | -1,37 | 19 | 5718912 | - | *LONP1* |
| cg03638795 | -5,06 | -254,32 | 114,77 | -9,20 | 11 | 416499 | + | *SIGIRR* |
| cg03643998 | 18,69 | -293,15 | 90,68 | 0,13 | 17 | 77042455 | - | *C1QTNF1* |
| cg03698343 | -12,00 | -315,12 | 187,58 | 4,52 | 9 | 139921892 | + | *ABCA2;C9orf13* |
| cg03725309 | -16,54 | -175,69 | 87,92 | -0,17 | 1 | 109757585 | + | *SARS* |
| cg03735592 | 1,66 | -245,01 | 213,08 | 6,99 | 6 | 138821354 | - | *NHSL1* |
| cg03746976 | -19,66 | -228,85 | 147,67 | 2,32 | 16 | 58035805 | - | *C16orf57* |
| cg03881294 | 28,20 | -216,31 | 88,92 | 1,39 | 2 | 11884333 | - | |
| cg03922748 | 11,92 | -241,06 | 79,62 | 2,09 | 2 | 220142903 | + | *DNAJB2* |
| cg03982897 | -20,37 | -292,90 | 198,44 | -3,57 | 11 | 1990108 | + | |
| cg04080625 | 7,46 | -268,41 | 146,41 | -4,91 | 1 | 15426634 | + | *KIAA1026* |
| cg04100595 | 5,63 | -150,16 | 75,64 | 5,24 | 12 | 27615138 | - | |
| cg04193015 | 4,93 | -207,20 | 176,67 | -2,65 | 17 | 35596680 | + | *ACACA* |
| cg04200607 | 3,66 | -207,63 | 107,88 | -2,85 | 10 | 72163475 | + | *EIF4EBP2* |
| cg04208403 | 3,20 | -318,53 | 236,09 | 5,06 | 16 | 49525807 | + | *ZNF423* |
| cg04308040 | 10,66 | -161,83 | 134,29 | 0,73 | 13 | 44942303 | + | |
| cg04424621 | -6,24 | -249,37 | 96,01 | 2,18 | 6 | 27101941 | + | *HIST1H2BJ* |

| | | | | | | | | |
|---|---|---|---|---|---|---|---|---|
| cg04425624 | 3,47 | -238,75 | 101,36 | 1,50 | 6 | 31543565 | - | *TNF* |
| cg04436528 | 23,19 | -313,55 | 237,13 | -0,90 | 8 | 143554875 | + | *BAI1* |
| cg04474832 | 1,87 | -318,26 | 142,24 | -1,35 | 3 | 52008487 | + | *ABHD14B;ABH* |
| cg04542977 | 14,58 | -185,62 | 106,88 | 3,86 | 1 | 51982669 | + | *EPS15* |
| cg04561237 | 9,95 | -247,07 | 87,52 | -3,91 | 10 | 126430405 | + | *FAM53B* |
| cg04581938 | -16,62 | -335,93 | 86,89 | 1,68 | 6 | 31364999 | - | |
| cg04596060 | -0,70 | -309,79 | 86,81 | 8,04 | 12 | 114404600 | + | *RBM19* |
| cg04604946 | -16,82 | -279,17 | 153,34 | 4,31 | 12 | 7023352 | + | *LRRC23;ENO2* |
| cg04651240 | -5,86 | -238,23 | 162,72 | 1,17 | 11 | 18229519 | - | *LOC494141* |
| cg04666029 | -10,46 | -419,15 | 101,85 | -1,58 | 11 | 2552843 | - | *KCNQ1* |
| cg04819580 | 1,87 | -255,71 | 106,45 | -3,16 | 7 | 150635526 | + | |
| cg04872610 | 17,99 | -303,70 | 140,74 | -8,12 | 1 | 114448489 | - | *AP4B1* |
| cg04875128 | 14,79 | 168,76 | 10,48 | 0,81 | 15 | 31775895 | - | *OTUD7A* |
| cg04890576 | 3,71 | -147,88 | 108,77 | 1,23 | 17 | 73032613 | + | |
| cg04956949 | 20,07 | -258,85 | 191,77 | 1,58 | 7 | 73119262 | + | *STX1A* |
| cg04959790 | 8,63 | -234,97 | 182,52 | 3,92 | 11 | 47278977 | + | *NR1H3* |
| cg04999352 | 1,72 | -272,81 | 109,02 | 4,92 | 11 | 63304614 | - | *RARRES3* |
| cg05045027 | 1,83 | -226,11 | 133,14 | -3,31 | 11 | 119331282 | + | |
| cg05061804 | -15,69 | -287,78 | 118,27 | -2,66 | 9 | 135282417 | + | *TTF1* |
| cg05091997 | 4,87 | -196,03 | 80,75 | -0,77 | 17 | 60897721 | + | |
| cg05156137 | -3,56 | -154,73 | 100,47 | -0,97 | 21 | 35898975 | - | *RCAN1* |
| cg05191655 | 4,41 | -238,15 | 109,75 | 0,92 | 4 | 38162793 | - | |
| cg05207048 | -5,78 | -147,46 | 79,61 | 0,62 | 5 | 167513456 | - | *ODZ2* |
| cg05237436 | -0,80 | -157,19 | 103,98 | -1,64 | 5 | 138533428 | + | *SIL1* |
| cg05242244 | -6,57 | -320,59 | 183,73 | 4,01 | 12 | 53553086 | - | *CSAD* |
| cg05308819 | -3,67 | -241,64 | 178,70 | -6,06 | 1 | 155959156 | - | |
| cg05316627 | 7,55 | -185,21 | 154,67 | -0,30 | 6 | 87861261 | - | |
| cg05324516 | -4,65 | -252,03 | 232,93 | 2,79 | 10 | 29421357 | - | |
| cg05369942 | 2,82 | -264,76 | 102,42 | -4,55 | 4 | 142228617 | - | |
| cg05379350 | -9,52 | -257,55 | 114,82 | -1,98 | 17 | 27917157 | - | *GIT1* |
| cg05405914 | -1,95 | -228,23 | 172,28 | 0,78 | 16 | 8972233 | + | |
| cg05412028 | -2,14 | -125,85 | 67,37 | -2,16 | 13 | 95952937 | - | *ABCC4* |
| cg05584950 | 3,76 | -212,69 | 93,92 | -2,25 | 2 | 86012847 | - | *ATOH8* |
| cg05619598 | -6,09 | -197,92 | 136,07 | -0,99 | 19 | 33569228 | - | |
| cg05694021 | 1,07 | -119,89 | 91,06 | -3,35 | 12 | 19699504 | + | |
| cg06007201 | -7,55 | -311,93 | 89,29 | -2,05 | 16 | 88850218 | + | *FAM38A* |
| cg06142740 | 5,91 | -300,47 | 138,22 | -1,01 | 2 | 28973577 | + | *PPP1CB* |
| cg06163904 | -0,21 | -297,36 | 240,34 | -0,87 | 17 | 78964779 | + | *CHMP6* |
| cg06240854 | -2,73 | -217,23 | 145,73 | 4,17 | 2 | 99771011 | - | *LIPT1* |
| cg06247837 | -4,13 | -201,15 | 168,69 | 2,28 | 17 | 37820135 | - | *TCAP* |
| cg06285727 | 35,75 | -213,65 | 86,43 | -6,26 | 11 | 72524028 | + | *ATG16L2* |
| cg06413398 | -11,54 | -137,38 | 97,89 | -0,39 | 6 | 110736865 | + | *DDO* |
| cg06437747 | -8,05 | -222,86 | 119,46 | -1,06 | 1 | 2890836 | - | |
| cg06484360 | -9,11 | -271,89 | 86,62 | -0,59 | 19 | 18434522 | - | *LSM4* |
| cg06567855 | -13,18 | -338,56 | 99,61 | -2,33 | 2 | 85977911 | - | |
| cg06639320 | 43,26 | 252,91 | -59,85 | 6,84 | 2 | 106015739 | - | *FHL2* |
| cg06647068 | -8,45 | -197,14 | 118,13 | 1,19 | 12 | 104853274 | - | *CHST11* |
| cg06661266 | 5,07 | -200,46 | 162,80 | -0,32 | 3 | 134027721 | - | |
| cg06685111 | -1,22 | -323,40 | 180,94 | 2,02 | 6 | 30295466 | - | *HCG18* |

| | | | | | | | | |
|---|---|---|---|---|---|---|---|---|
| cg06777902 | 4,17 | -190,67 | 96,93 | -3,86 | 10 | 127622996 | - | *FANK1* |
| cg06819357 | 0,68 | -231,21 | 198,71 | -3,14 | 14 | 102928437 | + | *TECPR2* |
| cg06911110 | 13,08 | -200,77 | 172,98 | -0,08 | 1 | 17293971 | - | *CROCC* |
| cg06934523 | -4,02 | -270,76 | 141,28 | -2,81 | 14 | 70077930 | - | *KIAA0247* |
| cg07027613 | -11,51 | -197,46 | 92,91 | 5,55 | 12 | 7260608 | - | *C1RL* |
| cg07080372 | -28,87 | -267,00 | 107,14 | 4,06 | 11 | 796607 | - | *SLC25A22* |
| cg07082267 | -19,67 | -209,06 | 142,41 | 2,05 | 16 | 85429035 | - | |
| cg07092212 | 14,79 | -213,04 | 87,87 | -0,03 | 11 | 46382544 | - | *DGKZ* |
| cg07127410 | -0,92 | -177,01 | 151,78 | -2,58 | 22 | 29427851 | - | *ZNRF3* |
| cg07164639 | -3,45 | -125,35 | 105,16 | -0,03 | 6 | 110736958 | + | *DDO* |
| cg07191657 | 4,39 | -189,52 | 137,76 | -0,04 | 10 | 14478541 | + | *MIR1265* |
| cg07211259 | -20,47 | -186,97 | 93,34 | -3,15 | 9 | 5510497 | - | *PDCD1LG2* |
| cg07234388 | 7,10 | -276,88 | 231,46 | -4,10 | 1 | 156261841 | + | *TMEM79* |
| cg07368443 | -6,85 | -295,40 | 166,98 | -0,73 | 1 | 45265337 | + | *PLK3* |
| cg07388493 | 0,73 | -192,70 | 144,03 | -2,32 | 1 | 39491459 | - | *NDUFS5* |
| cg07568841 | -6,12 | -191,56 | 103,13 | -1,15 | 7 | 30362781 | + | *ZNRF2* |
| cg07583137 | -11,28 | -205,73 | 192,32 | 0,60 | 8 | 82644012 | - | *CHMP4C* |
| cg07703358 | 5,38 | -259,01 | 95,15 | 3,29 | 7 | 138702117 | - | |
| cg07843120 | -17,70 | -264,09 | 86,28 | 1,03 | 19 | 34856957 | - | *GPI* |
| cg07850154 | 0,52 | -191,20 | 125,42 | 2,84 | 5 | 63461232 | + | *RNF180* |
| cg07931844 | -9,24 | -256,80 | 146,83 | -0,96 | 15 | 72102213 | - | *NR2E3* |
| cg08090640 | 0,75 | -219,26 | 157,99 | 4,61 | 17 | 41159289 | - | *IFI35* |
| cg08128734 | -8,44 | -156,31 | 138,19 | -2,66 | 1 | 206685423 | - | *RASSF5* |
| cg08138505 | -18,03 | -353,07 | 100,07 | 4,99 | 11 | 67118746 | - | *LOC100130987* |
| cg08160331 | 3,43 | 274,45 | -76,25 | -3,16 | 11 | 75140865 | + | *KLHL35* |
| cg08234504 | -13,07 | -279,76 | 137,63 | 0,68 | 5 | 139013317 | + | |
| cg08262002 | 4,14 | -184,95 | 151,48 | -0,48 | 4 | 16575323 | + | *LDB2* |
| cg08301612 | 4,68 | -175,99 | 109,32 | 0,66 | 17 | 43228571 | + | *HEXIM1* |
| cg08337633 | 22,78 | -385,83 | 83,83 | 5,51 | 7 | 55602109 | - | *VOPP1* |
| cg08343101 | 18,77 | -290,94 | 183,96 | 0,09 | 11 | 44933332 | - | *TSPAN18* |
| cg08409562 | 11,88 | -220,04 | 168,08 | 0,10 | 6 | 31737885 | - | *C6orf27* |
| cg08415592 | 2,45 | -228,55 | 113,12 | 6,45 | 22 | 36648973 | - | *APOL1* |
| cg08426733 | -12,73 | -306,81 | 168,88 | 0,63 | 1 | 1376740 | - | *VWA1* |
| cg08453194 | 9,26 | -314,30 | 180,23 | 0,94 | 6 | 41904398 | + | *CCND3* |
| cg08468689 | -3,99 | -254,36 | 113,24 | -1,37 | 17 | 40346680 | - | *GHDC* |
| cg08471846 | 20,47 | -267,28 | 132,19 | 0,96 | 19 | 799621 | - | *PTBP1* |
| cg08511485 | -1,25 | -252,56 | 90,62 | 1,30 | 3 | 168945731 | - | *MECOM* |
| cg08541155 | -5,90 | -188,63 | 113,25 | 0,16 | 3 | 118994052 | + | |
| cg08570034 | -3,07 | -166,10 | 141,42 | -2,35 | 5 | 175227702 | + | *CPLX2* |
| cg08644498 | 16,85 | -318,24 | 188,11 | -1,21 | 1 | 46502608 | + | |
| cg08713098 | -19,92 | -264,02 | 114,70 | -1,41 | 7 | 100005500 | - | *ZCWPW1* |
| cg08761208 | -12,68 | -309,43 | 119,55 | -5,14 | 15 | 65693289 | + | *IGDCC4* |
| cg08784091 | 20,57 | -436,82 | 108,76 | -2,01 | 6 | 31670005 | - | *BAT5* |
| cg08877357 | 2,23 | -136,48 | 94,70 | 2,80 | 10 | 113120532 | + | |
| cg08913523 | -3,98 | -189,17 | 93,39 | 1,39 | 8 | 126649807 | + | |
| cg08957484 | 12,33 | 198,18 | -25,13 | 6,49 | 5 | 132083532 | + | *CCNI2* |
| cg09124496 | -1,93 | -176,00 | 164,47 | 3,01 | 7 | 41735851 | + | *LOC285954* |
| cg09143195 | -11,29 | -162,82 | 88,94 | -4,66 | 12 | 41085901 | - | *CNTN1* |
| cg09294739 | -10,31 | -147,58 | 79,27 | 0,80 | 16 | 55218782 | - | |

| | | | | | | | | |
|---|---|---|---|---|---|---|---|---|
| cg09308553 | 1,56 | -280,58 | 157,71 | -1,63 | 16 | 56763573 | - | *NUP93* |
| cg09340639 | -2,50 | -231,57 | 84,57 | 2,12 | 1 | 157789662 | - | *FCRL1* |
| cg09608765 | 5,19 | -266,54 | 120,69 | 0,79 | 3 | 45636137 | + | *LIMD1* |
| cg09642020 | -7,43 | -311,17 | 201,35 | -1,71 | 9 | 139379173 | + | *C9orf163* |
| cg09748749 | -11,67 | -228,12 | 145,98 | 1,62 | 7 | 65540429 | + | *ASL* |
| cg09809672 | -11,12 | -163,69 | 131,16 | 3,87 | 1 | 236557682 | - | *EDARADD* |
| cg10052840 | -13,04 | -265,32 | 201,66 | -0,84 | 19 | 4558119 | - | *SEMA6B* |
| cg10137837 | 12,88 | 245,62 | -34,67 | 3,56 | 17 | 6926742 | + | *BCL6B* |
| cg10448052 | 12,43 | -223,96 | 90,50 | -4,44 | 12 | 53715787 | + | *AAAS* |
| cg10476085 | 19,04 | -259,86 | 175,31 | 1,33 | 5 | 142065737 | - | *FGF1* |
| cg10650821 | -6,72 | -342,75 | 105,94 | 0,68 | 6 | 31543686 | + | *TNF* |
| cg10717214 | 4,33 | -315,76 | 100,66 | 1,53 | 6 | 31543557 | - | *TNF* |
| cg10986043 | 6,41 | -226,08 | 171,09 | 0,97 | 17 | 37820495 | - | *TCAP* |
| cg11076306 | -9,99 | -193,78 | 177,06 | -4,53 | 4 | 41430667 | + | *LIMCH1* |
| cg11142333 | 7,79 | -275,46 | 239,90 | 3,32 | 1 | 173840261 | + | *ZBTB37* |
| cg11194994 | 0,72 | -296,60 | 134,06 | -3,59 | 1 | 160175974 | - | *PEA15* |
| cg11331344 | 7,19 | -288,68 | 143,33 | 0,50 | 17 | 73629463 | - | *RECQL5* |
| cg11344352 | -7,29 | -366,89 | 70,09 | 2,15 | 19 | 45927696 | + | *ERCC1* |
| cg11436113 | -4,31 | -234,05 | 189,57 | 2,44 | 20 | 19191145 | - | |
| cg11453058 | -2,97 | -260,29 | 104,34 | -2,18 | 19 | 14064176 | - | *PODNL1* |
| cg11619216 | -3,65 | -258,29 | 213,67 | 0,17 | 17 | 73642080 | - | *RECQL5* |
| cg11741201 | 12,82 | -210,59 | 102,49 | 0,69 | 11 | 35638398 | - | *FJX1* |
| cg11807280 | -2,14 | -120,85 | 111,53 | 0,26 | 2 | 66654644 | + | |
| cg11826475 | 13,59 | -299,07 | 156,00 | 1,68 | 12 | 120704006 | - | *PXN* |
| cg11836829 | -3,16 | -251,32 | 115,15 | 1,31 | 1 | 212605702 | - | *NENF* |
| cg12009872 | 7,26 | -269,47 | 113,12 | -2,21 | 15 | 51520739 | - | *CYP19A1* |
| cg12068124 | -4,41 | -289,10 | 104,00 | 4,15 | 19 | 1238538 | + | *C19orf26* |
| cg12079303 | -8,01 | -175,75 | 118,32 | 2,06 | 1 | 61547163 | - | *NFIA* |
| cg12179661 | 16,66 | -269,98 | 194,66 | 7,92 | 9 | 140333805 | + | *ENTPD8* |
| cg12197142 | 13,30 | -315,66 | 115,96 | -0,70 | 17 | 73629245 | - | *RECQL5;LOC[illegible]* |
| cg12261786 | -11,36 | -256,31 | 233,07 | -7,12 | 10 | 88727830 | + | *C10orf116* |
| cg12278474 | 13,28 | -206,55 | 117,51 | 0,51 | 1 | 5221357 | + | |
| cg12283460 | -0,15 | -247,74 | 94,11 | 1,71 | 15 | 59279556 | + | *RNF111* |
| cg12483947 | -14,22 | -208,51 | 111,56 | 0,41 | 10 | 72640100 | - | *SGPL1* |
| cg12526474 | 7,94 | -336,58 | 125,92 | 0,08 | 7 | 140097579 | - | *SLC37A3* |
| cg12554573 | -5,90 | -307,33 | 254,28 | -6,71 | 3 | 51976667 | - | *PARP3;RRP9* |
| cg12623930 | 5,33 | -324,35 | 129,20 | 0,80 | 3 | 52008802 | - | *ABHD14B;AB[illegible]* |
| cg12681001 | 12,46 | -322,11 | 95,77 | -5,15 | 6 | 31543540 | - | *TNF* |
| cg12688670 | 27,75 | -322,50 | 134,27 | -0,01 | 16 | 29801602 | + | *KIF22* |
| cg12706425 | 1,95 | -221,56 | 90,84 | -2,62 | 9 | 73029578 | + | *KLF9* |
| cg12899747 | -13,84 | -168,66 | 103,65 | 0,85 | 3 | 25391527 | - | |
| cg13033938 | -9,09 | -428,42 | 80,91 | 5,59 | 3 | 49824475 | - | *IP6K1* |
| cg13055199 | 4,94 | -203,19 | 147,42 | -7,21 | 15 | 75136278 | - | *ULK3* |
| cg13066481 | 16,36 | -228,44 | 161,87 | -3,76 | 3 | 123372199 | - | *MYLK* |
| cg13072940 | 8,69 | -256,79 | 143,29 | -3,27 | 3 | 49967521 | + | *MON1A* |
| cg13120986 | 1,71 | -189,15 | 100,79 | -4,51 | 12 | 7168012 | - | *C1S* |
| cg13203811 | 40,78 | -254,49 | 164,10 | -1,88 | 12 | 58136245 | - | *AGAP2* |
| cg13319938 | -25,50 | -321,39 | 257,70 | -6,83 | 11 | 2436251 | - | *TRPM5* |
| cg13385220 | 7,87 | -194,42 | 175,39 | -1,56 | 1 | 202250453 | + | *LGR6* |

| | | | | | | | | |
|---|---|---|---|---|---|---|---|---|
| cg13420364 | 16,80 | -157,69 | 120,15 | 0,34 | 1 | 234857659 | - | |
| cg13428009 | -13,91 | -371,17 | 89,29 | -1,74 | 9 | 139383171 | - | |
| cg13458211 | -2,82 | -263,78 | 172,52 | -5,73 | 1 | 44444217 | + | *B4GALT2* |
| cg13501527 | 12,33 | -292,84 | 211,66 | 3,66 | 9 | 123640474 | - | *PHF19* |
| cg13640414 | -0,33 | -203,94 | 131,41 | 2,75 | 19 | 15530870 | + | *AKAP8L* |
| cg13683374 | -3,27 | -235,36 | 138,93 | -0,44 | 17 | 72364767 | - | *GPR142* |
| cg13709639 | -3,68 | -204,32 | 83,91 | -1,88 | 12 | 49526040 | + | *TUBA1B* |
| cg13790426 | 13,23 | -219,60 | 82,78 | 1,34 | 1 | 109506963 | - | *CLCC1* |
| cg13807549 | -3,74 | -220,42 | 129,11 | 3,45 | 9 | 116444721 | - | |
| cg13823169 | 5,90 | -247,99 | 171,51 | 0,97 | 9 | 139776893 | - | |
| cg14039301 | -9,69 | -296,07 | 217,98 | 4,24 | 11 | 296540 | + | |
| cg14042143 | -2,02 | -191,94 | 183,64 | 0,92 | 7 | 2646782 | - | *IQCE* |
| cg14058848 | -17,14 | -258,49 | 202,96 | 1,20 | 9 | 130479114 | - | *TTC16;PTRH1* |
| cg14175438 | -8,66 | -260,71 | 107,65 | 8,26 | 7 | 121036729 | + | *FAM3C* |
| cg14188401 | -8,95 | -296,18 | 256,90 | 0,75 | 3 | 134097684 | - | |
| cg14292522 | 8,73 | -183,12 | 121,91 | 2,70 | 12 | 100535489 | + | *UHRF1BP1L* |
| cg14305711 | -3,85 | -225,88 | 88,10 | 1,42 | 2 | 62132430 | + | *COMMD1* |
| cg14314729 | 3,51 | -225,50 | 146,02 | -0,29 | 5 | 179815975 | - | |
| cg14359680 | -20,10 | -272,20 | 121,60 | 1,72 | 14 | 105532208 | - | *GPR132* |
| cg14583999 | -6,02 | -142,51 | 124,39 | 1,38 | 3 | 10019040 | - | *TMEM111* |
| cg14609289 | 7,37 | -318,36 | 226,19 | 0,66 | 17 | 77083037 | + | *ENGASE* |
| cg14671809 | 14,85 | -231,07 | 183,27 | 0,73 | 3 | 55558213 | - | *ERC2* |
| cg14775286 | 10,45 | -203,10 | 131,28 | 3,11 | 3 | 137833494 | + | *DZIP1L* |
| cg14837598 | -10,15 | -220,67 | 191,64 | 1,59 | 8 | 21916635 | - | *EPB49* |
| cg14898223 | -0,49 | -147,98 | 123,86 | 2,32 | 2 | 190447407 | + | |
| cg14934280 | 31,20 | -285,25 | 120,40 | 3,58 | 19 | 47494587 | + | *GRLF1* |
| cg14956327 | 9,60 | -134,42 | 96,87 | 1,67 | 6 | 110737053 | - | *DDO* |
| cg14963724 | -6,40 | -222,69 | 79,36 | 6,84 | 18 | 72166303 | - | *CNDP2* |
| cg14973055 | 11,24 | -176,27 | 138,41 | 5,80 | 17 | 72306038 | - | *DNAI2* |
| cg14977938 | 6,85 | -258,46 | 231,65 | -1,72 | 14 | 104190829 | + | *ZFYVE21* |
| cg14989226 | 15,51 | -219,92 | 118,18 | -0,31 | 7 | 75897850 | - | *SRRM3* |
| cg15034393 | 28,13 | -150,56 | 101,63 | 0,89 | 3 | 152856773 | - | |
| cg15125438 | -7,91 | -293,54 | 255,00 | -1,88 | 12 | 113684389 | + | *TPCN1* |
| cg15298486 | 10,19 | -274,12 | 99,10 | -5,76 | 10 | 102046690 | + | *BLOC1S2* |
| cg15393702 | 3,30 | -191,53 | 152,34 | -0,39 | 18 | 21243529 | + | *ANKRD29* |
| cg15416179 | -26,46 | -310,10 | 71,25 | 4,35 | 17 | 21189859 | - | *MAP2K3* |
| cg15538427 | 0,56 | -284,81 | 203,20 | -1,26 | 11 | 62457014 | - | *LRRN4CL* |
| cg15704699 | 13,31 | -230,64 | 203,01 | 3,72 | 17 | 26439377 | + | *NLK* |
| cg15743533 | 22,43 | -262,75 | 100,05 | 7,50 | 20 | 815345 | + | *FAM110A* |
| cg15804973 | 1,55 | -177,55 | 118,91 | -0,92 | 6 | 137114513 | + | *MAP3K5* |
| cg15829826 | 3,74 | -411,22 | 87,36 | 1,61 | 11 | 65153860 | + | *FRMD8* |
| cg15845821 | 16,65 | -192,41 | 141,59 | 0,81 | 19 | 16830613 | - | *NWD1* |
| cg15893346 | 3,66 | -224,32 | 131,55 | 2,16 | 7 | 65447565 | + | *GUSB* |
| cg15903032 | -6,50 | -228,96 | 161,30 | 2,26 | 10 | 101297605 | - | |
| cg16193278 | -7,16 | -264,85 | 103,98 | 0,54 | 13 | 100008450 | - | *UBAC2* |
| cg16273597 | -0,68 | -273,88 | 103,17 | 3,75 | 6 | 14117480 | + | *CD83* |
| cg16276108 | -5,36 | -226,19 | 143,76 | 0,19 | 5 | 180213189 | + | |
| cg16363586 | 7,30 | -241,65 | 91,83 | -0,51 | 19 | 17516329 | - | *BST2* |
| cg16541026 | -2,21 | -236,27 | 145,10 | 0,31 | 3 | 49026934 | - | *P4HTM* |

| | | | | | | | | |
|---|---|---|---|---|---|---|---|---|
| cg16618104 | 6,08 | -252,79 | 103,05 | -2,10 | 12 | 104853100 | + | *CHST11* |
| cg16640358 | -5,66 | -279,38 | 150,10 | 1,20 | 2 | 239892057 | - | |
| cg16664617 | -14,33 | -293,24 | 249,71 | -3,10 | 19 | 17607007 | - | *SLC27A1* |
| cg16677191 | -16,19 | -191,76 | 98,02 | -1,25 | 5 | 95159423 | - | *GLRX* |
| cg16742481 | -14,93 | -246,35 | 155,94 | -0,49 | 4 | 157090555 | - | |
| cg16744741 | -1,77 | -161,72 | 117,41 | 2,10 | 4 | 82126025 | - | *PRKG2* |
| cg16762684 | 17,79 | -198,51 | 69,02 | 3,01 | 18 | 74820493 | + | *MBP* |
| cg16810343 | 3,09 | -244,57 | 136,21 | -0,50 | 7 | 75898970 | + | *SRRM3* |
| cg16867657 | 48,86 | 173,46 | -55,16 | 5,22 | 6 | 11044877 | + | *ELOVL2* |
| cg16960758 | 26,38 | -198,22 | 165,68 | -0,99 | 2 | 169658992 | - | *NOSTRIN* |
| cg16983588 | -6,27 | -157,18 | 127,01 | -0,09 | 11 | 129793108 | + | *PRDM10* |
| cg17133388 | 13,00 | -357,00 | 98,64 | 1,74 | 3 | 122102727 | + | *FAM162A* |
| cg17168836 | -15,50 | -191,31 | 115,17 | 1,29 | 1 | 68256161 | - | *GNG12* |
| cg17183905 | -26,14 | -242,45 | 107,33 | -0,34 | 12 | 110253842 | - | *TRPV4* |
| cg17329534 | -1,71 | -228,74 | 99,00 | 0,04 | 1 | 154980743 | - | *ZBTB7B* |
| cg17436656 | 2,13 | -231,24 | 174,78 | -0,03 | 12 | 53627106 | + | *RARG* |
| cg17457912 | 5,23 | -228,33 | 161,74 | -6,58 | 17 | 1617102 | - | *C17orf91* |
| cg17593342 | 4,74 | -208,72 | 177,84 | -4,86 | 6 | 14037614 | - | |
| cg17621438 | -0,71 | -202,33 | 116,50 | -1,02 | 5 | 63461216 | + | *RNF180* |
| cg17721618 | -18,30 | -322,23 | 176,56 | 6,31 | 15 | 42376692 | - | *PLA2G4D* |
| cg18076651 | -9,90 | -347,32 | 132,87 | -1,46 | 12 | 53625605 | - | *RARG* |
| cg18079948 | 3,55 | -134,09 | 93,42 | -1,53 | 10 | 20009221 | - | |
| cg18150280 | 8,53 | -202,25 | 107,78 | 7,73 | 1 | 192776859 | - | *RGS2* |
| cg18186343 | -8,02 | -179,85 | 118,20 | 3,05 | 14 | 101317620 | - | *MIR770* |
| cg18215449 | -6,83 | -177,35 | 131,88 | 6,53 | 12 | 66089473 | + | |
| cg18333339 | 8,15 | -231,73 | 173,44 | -0,32 | 3 | 112885856 | - | |
| cg18450254 | -6,16 | -126,82 | 94,13 | -1,28 | 3 | 64200005 | + | *PRICKLE2* |
| cg18505959 | -21,46 | -387,82 | 112,64 | -0,35 | 6 | 31670195 | - | *BAT5* |
| cg18568843 | -13,99 | -353,69 | 110,82 | -0,94 | 6 | 52535550 | + | *TMEM14A* |
| cg18651026 | -10,21 | -270,92 | 203,20 | 2,32 | 6 | 33140660 | - | *COL11A2* |
| cg18715243 | -1,80 | -262,98 | 221,76 | 1,19 | 8 | 37658755 | - | *GPR124* |
| cg18738190 | 14,72 | -153,90 | 132,31 | -0,37 | 10 | 73740291 | + | *CHST3* |
| cg18779283 | -14,71 | -365,20 | 107,52 | 0,49 | 3 | 45636154 | + | *LIMD1* |
| cg18786171 | 9,15 | -248,96 | 157,65 | 1,85 | 10 | 75935758 | + | *ADK* |
| cg18797590 | 0,18 | -191,60 | 164,95 | 7,38 | 4 | 15480643 | - | *CC2D2A* |
| cg18808904 | -2,59 | -194,23 | 137,47 | -3,05 | 20 | 11898851 | - | *BTBD3* |
| cg18826637 | -9,64 | -138,02 | 123,75 | 1,89 | 2 | 145116633 | + | |
| cg18887458 | 3,91 | -114,39 | 85,51 | 6,05 | 7 | 115995934 | + | |
| cg18933331 | -6,38 | -210,85 | 151,07 | 2,60 | 1 | 110186418 | - | |
| cg18993949 | 5,35 | -263,76 | 203,05 | -0,01 | 4 | 15481269 | + | *CC2D2A* |
| cg19025497 | -9,54 | -284,05 | 97,12 | -0,79 | 5 | 138534324 | + | *SIL1* |
| cg19056004 | 10,84 | -227,46 | 173,68 | 0,52 | 12 | 7023262 | - | *LRRC23;ENO2* |
| cg19261426 | -2,63 | -209,97 | 171,24 | -2,16 | 1 | 228560439 | + | *OBSCN* |
| cg19270739 | 20,34 | -181,04 | 125,21 | -1,63 | 1 | 1368846 | - | |
| cg19344626 | -16,80 | -160,82 | 164,61 | 3,47 | 19 | 16830749 | - | *NWD1* |
| cg19381811 | -5,63 | -190,15 | 103,44 | 3,28 | 3 | 49851713 | - | *UBA7* |
| cg19433091 | 17,07 | -253,45 | 120,00 | 3,23 | 15 | 43426498 | - | *TMEM62* |
| cg19453093 | -0,07 | -197,53 | 140,02 | -0,75 | 14 | 88655985 | + | *KCNK10* |
| cg19500607 | -5,25 | -243,29 | 205,69 | 3,80 | 5 | 148034319 | + | *HTR4* |

| | | | | | | | | |
|---|---|---|---|---|---|---|---|---|
| cg19578183 | -7,97 | -265,46 | 207,95 | 0,10 | 2 | 128991530 | + | |
| cg19612068 | 23,19 | -248,98 | 158,25 | -0,63 | 6 | 33130024 | + | |
| cg19670290 | -7,57 | -227,09 | 135,21 | 1,24 | 15 | 91477210 | - | *HDDC3* |
| cg19722847 | -4,47 | -266,35 | 98,96 | 0,46 | 12 | 30849114 | + | *IPO8* |
| cg19729744 | 10,11 | -151,66 | 134,86 | -1,78 | 3 | 194752020 | + | |
| cg19753867 | 18,32 | -279,61 | 139,57 | -1,15 | 20 | 35383166 | - | *DSN1* |
| cg19758448 | -0,29 | -169,21 | 145,82 | -2,86 | 17 | 37828296 | + | *PGAP3* |
| cg19761273 | 20,53 | -261,03 | 123,42 | 0,83 | 17 | 80232096 | + | *CSNK1D* |
| cg19784428 | 13,77 | -185,42 | 179,50 | 3,66 | 19 | 16830746 | - | *NWD1* |
| cg19804488 | -3,56 | -269,52 | 187,29 | -6,29 | 17 | 73760363 | - | *GALK1* |
| cg19838043 | 14,44 | -211,09 | 190,40 | 0,53 | 14 | 104196038 | - | *ZFYVE21* |
| cg19936954 | 7,20 | -259,98 | 124,24 | -2,11 | 1 | 180148192 | - | *QSOX1* |
| cg20052760 | 3,07 | -210,40 | 109,64 | -1,06 | 6 | 10510789 | + | |
| cg20067719 | 0,58 | -198,86 | 110,13 | 0,55 | 17 | 48623772 | + | *SPATA20* |
| cg20133890 | -3,05 | -217,05 | 159,64 | 3,36 | 6 | 31680144 | - | *LY6G6E* |
| cg20153322 | -20,16 | -245,84 | 187,60 | 0,53 | 12 | 120703977 | - | *PXN* |
| cg20222376 | 0,99 | -191,10 | 89,01 | -1,63 | 19 | 15530606 | - | *AKAP8L* |
| cg20249566 | -10,57 | -182,93 | 176,09 | 3,70 | 19 | 16830739 | - | *NWD1* |
| cg20267101 | 2,65 | -199,50 | 142,41 | -6,42 | 3 | 113006232 | - | *BOC;WDR52* |
| cg20303331 | -10,33 | -234,48 | 157,21 | -0,66 | 17 | 36945575 | - | *PIP4K2B* |
| cg20515136 | 4,47 | -226,62 | 133,18 | -4,81 | 3 | 159707145 | - | *IL12A* |
| cg20543183 | -19,01 | -277,48 | 211,44 | 1,81 | 6 | 31648146 | + | *LY6G5C* |
| cg20595453 | -18,16 | -227,30 | 187,71 | -0,17 | 6 | 33219392 | - | *VPS52* |
| cg20608990 | -1,99 | -199,22 | 124,16 | 1,50 | 2 | 202097607 | + | *CASP8* |
| cg20669012 | -6,95 | -249,98 | 124,96 | -3,62 | 3 | 11102341 | + | |
| cg20747538 | -12,77 | -247,03 | 130,92 | 1,12 | 3 | 137838021 | + | |
| cg20813374 | 5,53 | -253,22 | 161,81 | -0,33 | 6 | 35657180 | - | *FKBP5* |
| cg20822990 | -12,75 | -270,87 | 143,67 | -1,78 | 1 | 17338766 | - | *ATP13A2* |
| cg20912205 | 4,02 | -265,50 | 121,18 | -2,05 | 3 | 50337305 | - | *NAT6;HYAL3* |
| cg20964856 | 4,24 | -271,65 | 182,64 | -1,20 | 19 | 41767669 | - | *HNRNPUL1* |
| cg20988565 | -8,00 | -171,10 | 142,18 | -0,85 | 8 | 106334333 | + | *ZFPM2* |
| cg21120249 | -8,51 | -233,89 | 191,87 | 2,33 | 9 | 139921971 | - | *C9orf139* |
| cg21139312 | 8,12 | 300,34 | -217,34 | 3,88 | 17 | 55663225 | - | *MSI2* |
| cg21186955 | -2,39 | -224,32 | 118,83 | 1,70 | 7 | 100729412 | - | *TRIM56* |
| cg21222743 | -6,84 | -307,87 | 88,93 | 0,64 | 6 | 31543545 | - | *TNF* |
| cg21280392 | -0,28 | -232,27 | 158,20 | -2,36 | 17 | 47304116 | + | *PHOSPHO1* |
| cg21322248 | -55,50 | -376,28 | 85,78 | 0,52 | 15 | 77289047 | - | *PSTPIP1* |
| cg21333674 | -3,48 | -138,10 | 112,71 | 1,00 | 8 | 96705765 | - | |
| cg21406967 | -8,98 | -235,14 | 141,98 | 0,87 | 7 | 100464553 | - | *TRIP6* |
| cg21469505 | 10,81 | -195,72 | 122,74 | -0,06 | 18 | 47085749 | - | |
| cg21523751 | 2,02 | -180,70 | 94,77 | -0,30 | 1 | 182988639 | + | |
| cg21524899 | 9,29 | -325,95 | 279,38 | -0,27 | 1 | 225662606 | - | |
| cg21554217 | 1,58 | -295,31 | 107,96 | -0,47 | 5 | 138897467 | - | |
| cg21572722 | -9,77 | 333,99 | -87,27 | 6,39 | 6 | 11044894 | + | *ELOVL2* |
| cg21635307 | -10,63 | -268,86 | 105,08 | -2,04 | 6 | 90319877 | + | *ANKRD6* |
| cg21868031 | -33,79 | -387,58 | 90,18 | 1,09 | 1 | 207925112 | - | *CD46* |
| cg21878650 | -3,96 | -110,20 | 82,54 | 0,43 | 5 | 64558623 | + | *ADAMTS6* |
| cg21922223 | -8,72 | -343,10 | 106,99 | 3,90 | 17 | 75539118 | + | |
| cg21940640 | 5,65 | -210,38 | 173,54 | -0,67 | 7 | 56160893 | + | *PHKG1* |

| | | | | | | | | |
|---|---|---|---|---|---|---|---|---|
| cg21962791 | 1,17 | -204,91 | 134,47 | -2,33 | 12 | 21590305 | - | *PYROXD1* |
| cg21990700 | -2,63 | -207,22 | 108,02 | -1,91 | 12 | 7260776 | - | *LOC283314* |
| cg22156456 | 17,40 | -247,48 | 108,72 | 1,54 | 17 | 39844239 | - | *EIF1* |
| cg22156842 | -1,61 | -172,15 | 101,14 | 0,72 | 3 | 136537169 | - | *TMEM22* |
| cg22273555 | 1,86 | -231,68 | 137,81 | 5,00 | 6 | 33130034 | + | |
| cg22299097 | -11,98 | -203,49 | 113,38 | 1,86 | 3 | 97690590 | - | *MINA* |
| cg22361181 | -19,18 | -290,19 | 174,24 | -3,05 | 17 | 40171740 | - | *NKIRAS2* |
| cg22379463 | -15,89 | -253,37 | 192,09 | -1,92 | 3 | 45260330 | - | |
| cg22454769 | 1,25 | 149,85 | -29,55 | 8,85 | 2 | 106015767 | - | *FHL2* |
| cg22584802 | 3,55 | -214,53 | 91,14 | -3,27 | 7 | 129007902 | - | *AHCYL2* |
| cg22730004 | -3,80 | -140,52 | 93,22 | 1,84 | 1 | 158656718 | - | *SPTA1* |
| cg22737154 | -25,64 | -151,96 | 123,69 | -1,97 | 2 | 64631614 | - | |
| cg22747507 | 8,89 | -235,50 | 161,36 | 4,83 | 4 | 175635801 | - | *GLRA3* |
| cg22768222 | 5,48 | -146,75 | 111,32 | -9,10 | 6 | 45383690 | + | *RUNX2* |
| cg22864266 | 8,16 | -354,17 | 115,85 | 1,30 | 12 | 49319673 | - | *FKBP11* |
| cg22927302 | -27,38 | -312,35 | 222,58 | 3,02 | 3 | 50304463 | - | *SEMA3B* |
| cg22929506 | -10,00 | -214,65 | 123,43 | 2,14 | 2 | 219137490 | - | *PNKD* |
| cg22947000 | 9,74 | -171,18 | 118,64 | 0,59 | 16 | 81272281 | - | *BCMO1* |
| cg22976533 | 3,86 | -189,14 | 139,90 | 0,78 | 14 | 105784090 | - | *PACS2* |
| cg23003085 | -0,25 | -228,96 | 93,30 | -4,20 | 11 | 64084321 | - | *PRDX5* |
| cg23078123 | -13,92 | -232,69 | 187,66 | 7,08 | 1 | 68577796 | + | *GPR177* |
| cg23124451 | 8,67 | -268,34 | 184,82 | 1,64 | 22 | 39548131 | + | *CBX7* |
| cg23149300 | 0,71 | -239,89 | 125,37 | 0,40 | 10 | 36958940 | + | |
| cg23149687 | 8,36 | -166,55 | 90,20 | -5,17 | 5 | 119801643 | + | *PRR16* |
| cg23320649 | 1,58 | -255,59 | 185,01 | 0,56 | 3 | 50604613 | + | *C3orf18* |
| cg23341182 | -11,30 | -198,43 | 117,26 | 2,06 | 10 | 102046768 | + | *BLOC1S2* |
| cg23368715 | 1,06 | -222,64 | 153,53 | 3,01 | 12 | 7245510 | - | *C1R* |
| cg23389651 | 4,65 | -334,15 | 107,12 | 0,11 | 13 | 95327390 | + | |
| cg23460961 | -4,53 | -163,74 | 95,77 | -1,51 | 11 | 94382347 | + | |
| cg23461714 | -22,42 | -293,77 | 109,35 | -0,74 | 11 | 113184990 | - | *TTC12* |
| cg23677833 | -17,05 | -261,23 | 95,68 | 0,21 | 19 | 16308345 | - | *AP1M1* |
| cg23715749 | 14,85 | -187,19 | 130,80 | -1,53 | 1 | 37413867 | + | *GRIK3* |
| cg23718736 | -2,67 | -217,04 | 166,91 | 1,47 | 18 | 6413908 | - | *L3MBTL4* |
| cg23732483 | -5,71 | -163,08 | 87,48 | 1,41 | 3 | 48965611 | + | *ARIH2* |
| cg23744638 | -12,12 | -177,22 | 173,67 | 1,61 | 11 | 10323902 | - | |
| cg23836737 | 5,97 | -237,23 | 135,86 | 1,07 | 4 | 53411703 | + | |
| cg23950157 | -15,21 | -288,76 | 115,13 | -6,13 | 17 | 48275919 | + | *COL1A1* |
| cg23972551 | -9,52 | -360,46 | 102,49 | 1,05 | 11 | 68151664 | - | *LRP5* |
| cg24057710 | -1,55 | -253,92 | 184,59 | 2,76 | 11 | 126311095 | + | *KIRREL3* |
| cg24079702 | -1,04 | 197,30 | -21,96 | 7,32 | 2 | 106015771 | - | *FHL2* |
| cg24155190 | 0,12 | -303,80 | 89,83 | -0,80 | 1 | 201476619 | - | *CSRP1* |
| cg24711336 | 4,67 | -254,99 | 198,51 | 4,19 | 10 | 80063791 | - | |
| cg24774812 | -8,42 | -280,50 | 102,77 | 1,96 | 14 | 63743552 | - | *RHOJ* |
| cg24847230 | 29,00 | -345,09 | 194,89 | 1,35 | 17 | 46986807 | - | *UBE2Z* |
| cg24848615 | -4,31 | -220,80 | 187,81 | 0,61 | 19 | 3368396 | - | *NFIC* |
| cg24883498 | 0,81 | -264,05 | 120,12 | -0,54 | 19 | 18264771 | + | *PIK3R2* |
| cg24892069 | 1,73 | -122,69 | 118,06 | 4,72 | 10 | 33562205 | - | *NRP1* |
| cg24987259 | -0,29 | -223,14 | 103,71 | 3,30 | 11 | 66336293 | - | *CTSF* |
| cg25150953 | 3,41 | -126,31 | 77,61 | 1,53 | 4 | 41540229 | - | *LIMCH1* |

| | | | | | | | | |
|---|---|---|---|---|---|---|---|---|
| cg25268718 | 14,86 | -271,51 | 203,37 | -1,91 | 14 | 24604711 | + | *PSME1* |
| cg25311470 | 4,51 | -233,58 | 122,80 | -4,02 | 7 | 107950866 | + | *NRCAM* |
| cg25371036 | -3,41 | -231,10 | 202,60 | -0,16 | 11 | 94500749 | - | *AMOTL1* |
| cg25395188 | 9,46 | -251,24 | 95,73 | 0,12 | 11 | 60897782 | + | *VPS37C* |
| cg25413977 | 2,33 | -145,21 | 127,85 | -8,32 | 2 | 66651619 | + | |
| cg25424279 | 0,23 | -260,30 | 233,41 | 0,14 | 11 | 65683543 | + | |
| cg25439632 | 4,21 | -140,47 | 86,99 | 3,08 | 8 | 49892209 | + | |
| cg25616535 | -5,06 | -294,17 | 185,38 | 0,34 | 17 | 47294967 | - | *ABI3* |
| cg25782440 | -17,64 | -334,88 | 238,90 | -1,23 | 19 | 7979022 | - | *MAP2K7* |
| cg25793051 | 2,45 | -229,49 | 112,53 | 2,30 | 7 | 149439005 | - | |
| cg25994988 | -13,38 | -237,33 | 166,82 | 1,40 | 11 | 122652382 | + | *UBASH3B* |
| cg25998745 | 11,51 | -217,03 | 190,09 | 1,77 | 8 | 142028625 | + | |
| cg26094232 | 11,09 | -228,96 | 132,93 | -1,30 | 11 | 72148947 | + | |
| cg26101277 | -17,16 | -202,69 | 89,03 | -1,13 | 17 | 32690412 | - | *CCL1* |
| cg26158023 | -8,76 | -326,01 | 128,46 | 3,01 | 3 | 42881414 | + | *CCBP2* |
| cg26166595 | 3,08 | -211,39 | 128,53 | 2,78 | 11 | 11998715 | + | *DKK3* |
| cg26210267 | -3,56 | -303,00 | 118,61 | 1,68 | 4 | 668877 | + | *ATP5I* |
| cg26276120 | 4,90 | -312,80 | 116,93 | 5,26 | 12 | 6977747 | - | *TPI1* |
| cg26290219 | -12,78 | -216,50 | 156,60 | -3,10 | 6 | 33128906 | - | |
| cg26316599 | 8,43 | -207,54 | 154,55 | -2,14 | 5 | 172456055 | + | *ATP6V0E1* |
| cg26373518 | 8,52 | -216,11 | 138,73 | -3,12 | 22 | 31518942 | - | *INPP5J* |
| cg26450750 | -5,18 | -271,34 | 107,17 | -2,32 | 4 | 87816983 | - | |
| cg26483332 | -14,07 | -348,01 | 115,43 | -1,60 | 10 | 76993756 | + | *COMTD1* |
| cg26543112 | -12,35 | -234,51 | 107,03 | 0,34 | 6 | 133188277 | + | |
| cg26608718 | -8,45 | -153,44 | 100,70 | -1,62 | 19 | 15530737 | - | *AKAP8L* |
| cg26610808 | -20,20 | -291,32 | 100,24 | 1,58 | 10 | 102046685 | + | *BLOC1S2* |
| cg26614073 | -5,59 | -211,36 | 152,16 | 2,72 | 3 | 47517819 | - | *SCAP* |
| cg26748477 | 0,81 | -252,00 | 206,70 | 0,46 | 17 | 38516415 | + | |
| cg26787199 | 15,94 | -359,48 | 101,35 | -0,42 | 16 | 2044042 | - | *SYNGR3* |
| cg26808293 | -0,47 | -434,47 | 132,06 | -4,27 | 16 | 3072207 | - | *TNFRSF12A* |
| cg26894354 | -0,65 | -228,87 | 143,90 | -1,40 | 1 | 203311314 | - | *FMOD* |
| cg26954174 | 1,19 | -168,23 | 87,32 | 0,85 | 16 | 50730813 | - | *NOD2* |
| cg26963632 | 4,50 | -196,16 | 103,48 | -0,36 | 16 | 85558148 | - | |
| cg26969888 | 0,66 | -201,74 | 150,42 | -0,06 | 19 | 14064254 | - | *PODNL1* |
| cg27004870 | 10,10 | -328,68 | 100,33 | 3,05 | 16 | 88850384 | + | *FAM38A* |
| cg27209729 | -6,92 | -172,72 | 143,58 | -0,40 | 11 | 64428925 | + | *NRXN2* |
| cg27236973 | 2,57 | -160,22 | 95,13 | -4,11 | 17 | 39781997 | + | *KRT17* |
| cg27259408 | -2,44 | -258,10 | 183,70 | 3,73 | 19 | 10427154 | + | *FDX1L* |
| cg27269561 | -11,57 | -339,78 | 132,23 | -2,48 | 16 | 3072713 | + | *HCFC1R1* |
| cg27346545 | -5,29 | -200,51 | 132,47 | -7,38 | 20 | 1205378 | - | *RAD21L1* |
| cg27386529 | 3,29 | -227,37 | 142,67 | 0,70 | 3 | 47517807 | - | *SCAP* |
| cg27401724 | -7,04 | -272,41 | 215,54 | -3,82 | 17 | 43213629 | - | *ACBD4* |
| cg27409484 | 1,61 | -261,72 | 96,08 | -2,41 | 6 | 71721893 | + | |
| cg27470213 | 13,46 | -264,14 | 91,01 | 1,35 | 17 | 76967695 | - | *LGALS3BP* |
| ch.1.17167261 | -38,90 | -347,67 | 81,10 | -3,68 | 1 | 173405989 | + | |
| ch.1.839062R | -5,65 | -333,21 | 82,11 | 0,72 | 1 | 25282539 | + | *RUNX3* |
| ch.13.3956490 | 22,73 | -308,24 | 77,43 | 1,80 | 13 | 40666907 | + | |
| ch.14.9733109 | 72,73 | -394,17 | 71,25 | -3,05 | 14 | 98261346 | + | |
| ch.15.6779758 | -10,47 | -236,06 | 84,19 | 0,60 | 15 | 70010530 | + | |

| | | | | | | | |
|---|---|---|---|---|---|---|---|
| ch.19.1625111 | -12,63 | -346,33 | 71,26 | -0,69 | 19 | 16390119 | + |
| ch.2.10590135 | -0,63 | -233,23 | 72,08 | 0,39 | 2 | 106534922 | + |
| ch.2.20781454 | 2,99 | -257,09 | 71,31 | 1,39 | 2 | 208106299 | + |
| ch.2.217478R | -4,18 | -367,81 | 68,03 | 2,12 | 2 | 8142529 | + |
| ch.2.30415474 | -22,63 | -289,72 | 77,83 | -1,34 | 2 | 30561970 | + |
| ch.2.47286786 | 6,34 | -448,44 | 71,05 | 0,65 | 2 | 47433282 | + |
| ch.6.33611621 | -9,61 | -328,25 | 75,08 | 0,34 | 6 | 33503643 | + |

For each of the 491 methylation sites, we include the coefficients for the multivariate model, the slopes and intercepts for the average of n-independent linear regressions, and the weights for the WKDE model.

**Supplemental Table S4. Mortality-associated CpGs of the 491 CpG signature in LBC1921.**

| Site | HR | low CI 95% | up CI 95% | Pr(>\|z\|) | Slope (DNAm/age) | Gene | CHR |
|---|---|---|---|---|---|---|---|
| cg04875128 | 0,98 | 0,97 | 1,00 | 0,02 | 168,76 | *OTUD7A* | 15 |
| cg08128734 | 0,99 | 0,98 | 1,00 | 0,00 | -156,31 | *RASSF5* | 1 |
| cg11436113 | 0,98 | 0,97 | 0,99 | 0,01 | -234,05 | | 20 |
| cg19761273 | 0,97 | 0,95 | 1,00 | 0,04 | -261,03 | *CSNK1D* | 17 |
| cg19670290 | 0,97 | 0,96 | 0,98 | 0,00 | -227,09 | *HDDC3;UNC45A* | 15 |
| cg23003085 | 0,91 | 0,86 | 0,96 | 0,00 | -228,96 | *PRDX5;TRMT112* | 11 |
| cg23677833 | 0,89 | 0,84 | 0,95 | 0,00 | -261,23 | *AP1M1* | 19 |
| cg01812894 | 0,98 | 0,97 | 0,99 | 0,00 | -203,72 | *ALDH1A1* | 9 |
| cg23950157 | 0,94 | 0,91 | 0,97 | 0,00 | -288,76 | *COL1A1* | 17 |
| cg21990700 | 0,96 | 0,94 | 0,98 | 0,00 | -207,22 | *LOC283314;C1RL* | 12 |
| cg14934280 | 0,95 | 0,92 | 0,98 | 0,00 | -285,25 | *GRLF1* | 19 |
| cg06007201 | 0,91 | 0,86 | 0,96 | 0,00 | -311,93 | *FAM38A* | 16 |
| cg21635307 | 0,95 | 0,92 | 0,98 | 0,00 | -268,86 | *ANKRD6* | 6 |
| cg03206537 | 0,96 | 0,94 | 0,98 | 0,00 | -199,68 | *CTSA* | 20 |
| cg20813374 | 0,97 | 0,96 | 0,99 | 0,00 | -253,22 | *FKBP5* | 6 |
| cg27209729 | 0,98 | 0,97 | 0,99 | 0,00 | -172,72 | *NRXN2* | 11 |
| cg15393702 | 0,99 | 0,98 | 1,00 | 0,00 | -191,53 | *ANKRD29* | 18 |
| cg24155190 | 0,90 | 0,84 | 0,97 | 0,00 | -303,80 | *CSRP1* | 1 |
| cg17593342 | 0,98 | 0,97 | 1,00 | 0,00 | -208,72 | | 6 |
| cg09608765 | 0,96 | 0,94 | 0,99 | 0,00 | -266,54 | *LIMD1* | 3 |
| cg08913523 | 0,96 | 0,94 | 0,99 | 0,00 | -189,17 | | 8 |
| cg21922223 | 0,92 | 0,86 | 0,97 | 0,01 | -343,10 | | 17 |
| cg26158023 | 0,96 | 0,93 | 0,99 | 0,01 | -326,01 | *CCBP2* | 3 |
| cg19453093 | 0,98 | 0,97 | 1,00 | 0,01 | -197,53 | *KCNK1* | 14 |
| cg26808293 | 0,94 | 0,90 | 0,98 | 0,01 | -434,47 | *TNFRSF12A* | 16 |
| cg01243823 | 0,97 | 0,96 | 0,99 | 0,01 | -161,72 | *NOD2* | 16 |
| cg00602811 | 0,98 | 0,97 | 1,00 | 0,01 | -141,70 | *ZEB2* | 2 |
| ch.1.171672612F | 0,72 | 0,57 | 0,91 | 0,01 | -347,67 | | 1 |
| cg13823169 | 0,98 | 0,97 | 1,00 | 0,01 | -247,99 | | 9 |
| cg08301612 | 0,98 | 0,96 | 0,99 | 0,01 | -175,99 | *HEXIM1* | 17 |
| cg25994988 | 0,98 | 0,97 | 1,00 | 0,01 | -237,33 | *UBASH3B* | 11 |
| cg04080625 | 0,98 | 0,96 | 0,99 | 0,01 | -268,41 | *KIAA126* | 1 |
| cg18505959 | 0,93 | 0,87 | 0,98 | 0,01 | -387,82 | *BAT5* | 6 |
| cg06285727 | 0,94 | 0,90 | 0,99 | 0,01 | -213,65 | *ATG16L2* | 11 |
| cg21469505 | 0,98 | 0,97 | 1,00 | 0,01 | -195,72 | | 18 |
| cg03881294 | 0,95 | 0,92 | 0,99 | 0,01 | -216,31 | | 2 |
| cg04542977 | 0,98 | 0,97 | 1,00 | 0,01 | -185,62 | *EPS15* | 1 |
| cg25424279 | 0,97 | 0,96 | 0,99 | 0,01 | -260,30 | | 11 |
| cg27386529 | 0,98 | 0,97 | 1,00 | 0,01 | -227,37 | *SCAP* | 3 |
| cg26101277 | 0,96 | 0,93 | 0,99 | 0,01 | -202,69 | *CCL1* | 17 |
| cg05316627 | 0,99 | 0,98 | 1,00 | 0,01 | -185,21 | | 6 |
| cg25311470 | 0,98 | 0,96 | 0,99 | 0,01 | -233,58 | *NRCAM* | 7 |
| cg18797590 | 0,99 | 0,98 | 1,00 | 0,01 | -191,60 | *CC2D2A* | 4 |
| cg08471846 | 0,98 | 0,96 | 1,00 | 0,01 | -267,28 | *PTBP1* | 19 |
| cg14058848 | 0,98 | 0,96 | 1,00 | 0,01 | -258,49 | *TTC16;PTRH1* | 9 |

| | | | | | | | |
|---|---|---|---|---|---|---|---|
| cg12197142 | 0,94 | 0,90 | 0,99 | 0,01 | -315,66 | *RECQL5;LOC6438* | 17 |
| ch.2.47286786F | 0,34 | 0,15 | 0,81 | 0,01 | -448,44 | | 2 |
| cg19784428 | 0,99 | 0,98 | 1,00 | 0,02 | -185,42 | *NWD1* | 19 |
| cg04474832 | 0,97 | 0,95 | 0,99 | 0,02 | -318,26 | *ABHD14B;ABHD14A* | 3 |
| cg05379350 | 0,97 | 0,94 | 0,99 | 0,02 | -257,55 | *GIT1* | 17 |
| cg02610723 | 0,96 | 0,92 | 0,99 | 0,02 | -330,23 | *FAM38A* | 16 |
| cg03746976 | 0,98 | 0,96 | 1,00 | 0,02 | -228,85 | *C16orf57* | 16 |
| cg23836737 | 0,98 | 0,97 | 1,00 | 0,02 | -237,23 | | 4 |
| cg16677191 | 0,97 | 0,94 | 0,99 | 0,02 | -191,76 | *GLRX* | 5 |
| cg27470213 | 0,96 | 0,93 | 0,99 | 0,02 | -264,14 | *LGALS3BP* | 17 |
| cg12483947 | 0,97 | 0,95 | 1,00 | 0,02 | -208,51 | *SGPL1* | 10 |
| cg13072940 | 0,98 | 0,96 | 1,00 | 0,02 | -256,79 | *MON1A* | 3 |
| cg01955153 | 0,96 | 0,93 | 0,99 | 0,02 | -271,17 | | 16 |
| cg19729744 | 0,99 | 0,98 | 1,00 | 0,02 | -151,66 | | 3 |
| cg26894354 | 0,98 | 0,97 | 1,00 | 0,02 | -228,87 | *FMOD* | 1 |
| cg16363586 | 0,95 | 0,92 | 0,99 | 0,03 | -241,65 | *BST2* | 19 |
| cg14314729 | 0,98 | 0,97 | 1,00 | 0,03 | -225,50 | | 5 |
| cg07027613 | 0,96 | 0,92 | 1,00 | 0,03 | -197,46 | *C1RL;LOC283314* | 12 |
| ch.13.39564907R | 0,74 | 0,56 | 0,97 | 0,03 | -308,24 | | 13 |
| cg01511567 | 0,97 | 0,94 | 1,00 | 0,03 | -267,54 | *SSRP1* | 11 |
| cg18779283 | 0,93 | 0,87 | 0,99 | 0,03 | -365,20 | *LIMD1* | 3 |
| cg06647068 | 0,98 | 0,97 | 1,00 | 0,04 | -197,14 | *CHST11* | 12 |
| cg08090640 | 0,99 | 0,97 | 1,00 | 0,04 | -219,26 | *IFI35* | 17 |
| cg22864266 | 0,96 | 0,93 | 1,00 | 0,04 | -354,17 | *FKBP11* | 12 |
| cg13428009 | 0,86 | 0,75 | 0,99 | 0,04 | -371,17 | | 9 |
| cg05242244 | 0,98 | 0,97 | 1,00 | 0,04 | -320,59 | *CSAD* | 12 |
| cg19936954 | 0,98 | 0,95 | 1,00 | 0,04 | -259,98 | *QSOX1* | 1 |
| cg27004870 | 0,96 | 0,93 | 1,00 | 0,04 | -328,68 | *FAM38A* | 16 |
| cg16618104 | 0,96 | 0,93 | 1,00 | 0,04 | -252,79 | *CHST11* | 12 |
| cg26954174 | 0,97 | 0,95 | 1,00 | 0,04 | -168,23 | *NOD2* | 16 |
| cg22976533 | 0,99 | 0,98 | 1,00 | 0,04 | -189,14 | *PACS2* | 14 |
| cg08234504 | 0,97 | 0,95 | 1,00 | 0,04 | -279,76 | | 5 |
| cg20595453 | 0,99 | 0,97 | 1,00 | 0,04 | -227,30 | *VPS52* | 6 |
| cg03725309 | 0,96 | 0,92 | 1,00 | 0,04 | -175,69 | *SARS* | 1 |
| cg20747538 | 0,98 | 0,96 | 1,00 | 0,04 | -247,03 | | 3 |
| cg12179661 | 0,99 | 0,97 | 1,00 | 0,05 | -269,98 | *ENTPD8* | 9 |
| cg12009872 | 0,97 | 0,95 | 1,00 | 0,05 | -269,47 | *CYP19A1* | 15 |

The first 4 CpG sites are also mortality associated in the list of 27CpGs (with $R^2 > 0.7$ between DNAm and age in the training set).

**Supplemental Table S5. Mortality-associated CpGs of the 491 CpG signature in LBC1936.**

| Site | HR | low CI 95% | up CI 95% | Pr(>\|z\|) | Slope (DNAm/age) | Gene | CHR |
|---|---|---|---|---|---|---|---|
| cg11436113 | 0,97 | 0,95 | 0,98 | 0,00 | -234,05 | | 20 |
| cg01594949 | 0,97 | 0,96 | 0,99 | 0,00 | -239,86 | *NRARP* | 9 |
| cg05405914 | 0,97 | 0,96 | 0,99 | 0,00 | -228,23 | | 16 |
| cg14775286 | 0,97 | 0,96 | 0,99 | 0,00 | -203,10 | *DZIP1L* | 3 |
| cg14989226 | 0,97 | 0,95 | 0,99 | 0,00 | -219,92 | *SRRM3* | 7 |
| cg19578183 | 0,96 | 0,94 | 0,98 | 0,00 | -265,46 | | 2 |
| cg20813374 | 0,97 | 0,95 | 0,98 | 0,00 | -253,22 | *FKBP5* | 6 |
| cg01243823 | 0,97 | 0,96 | 0,99 | 0,00 | -161,72 | *NOD2* | 16 |
| cg06647068 | 0,98 | 0,96 | 0,99 | 0,00 | -197,14 | *CHST11* | 12 |
| cg08471846 | 0,96 | 0,94 | 0,99 | 0,00 | -267,28 | *PTBP1* | 19 |
| cg16541026 | 0,97 | 0,96 | 0,99 | 0,00 | -236,27 | *P4HTM* | 3 |
| cg16664617 | 0,96 | 0,94 | 0,98 | 0,00 | -293,24 | *SLC27A1* | 19 |
| cg17436656 | 0,98 | 0,96 | 0,99 | 0,00 | -231,24 | *RARG* | 12 |
| cg20595453 | 0,97 | 0,95 | 0,99 | 0,00 | -227,30 | *VPS52* | 6 |
| cg24848615 | 0,97 | 0,96 | 0,99 | 0,00 | -220,80 | *NFIC* | 19 |
| cg08713098 | 0,96 | 0,94 | 0,99 | 0,00 | -264,02 | *ZCWPW1* | 7 |
| cg20988565 | 0,98 | 0,97 | 0,99 | 0,00 | -171,10 | *ZFPM2* | 8 |
| cg23003085 | 0,92 | 0,88 | 0,97 | 0,00 | -228,96 | *PRDX5;TRMT112* | 11 |
| cg23677833 | 0,90 | 0,85 | 0,96 | 0,00 | -261,23 | *AP1M1* | 19 |
| cg11331344 | 0,97 | 0,94 | 0,99 | 0,00 | -288,68 | *RECQL5;LOC6438* | 17 |
| cg13501527 | 0,98 | 0,96 | 0,99 | 0,00 | -292,84 | *PHF19* | 9 |
| cg13823169 | 0,98 | 0,96 | 0,99 | 0,00 | -247,99 | | 9 |
| cg20912205 | 0,95 | 0,93 | 0,98 | 0,00 | -265,50 | *NAT6;HYAL3* | 3 |
| cg21990700 | 0,97 | 0,94 | 0,99 | 0,00 | -207,22 | *LOC283314;C1RL* | 12 |
| cg22156842 | 0,97 | 0,96 | 0,99 | 0,00 | -172,15 | *TMEM22* | 3 |
| cg05316627 | 0,99 | 0,97 | 1,00 | 0,00 | -185,21 | | 6 |
| cg14188401 | 0,96 | 0,94 | 0,99 | 0,00 | -296,18 | | 3 |
| cg19453093 | 0,98 | 0,97 | 0,99 | 0,00 | -197,53 | *KCNK1* | 14 |
| cg01234420 | 0,98 | 0,97 | 1,00 | 0,01 | -154,57 | *LOC15381* | 22 |
| cg02402091 | 0,98 | 0,96 | 0,99 | 0,01 | -218,54 | *ACSL5* | 10 |
| cg04581938 | 0,83 | 0,73 | 0,95 | 0,01 | -335,93 | | 6 |
| cg04959790 | 0,98 | 0,96 | 0,99 | 0,01 | -234,97 | *NR1H3* | 11 |
| cg05237436 | 0,98 | 0,96 | 0,99 | 0,01 | -157,19 | *SIL1* | 5 |
| cg19261426 | 0,98 | 0,97 | 1,00 | 0,01 | -209,97 | *OBSCN* | 1 |
| cg21139312 | 1,11 | 1,03 | 1,20 | 0,01 | 300,34 | *MSI2* | 17 |
| cg06240854 | 0,98 | 0,97 | 1,00 | 0,01 | -217,23 | *LIPT1;TSGA1* | 2 |
| cg08468689 | 0,97 | 0,95 | 0,99 | 0,01 | -254,36 | *GHDC* | 17 |
| cg24711336 | 0,98 | 0,96 | 0,99 | 0,01 | -254,99 | | 10 |
| cg26954174 | 0,97 | 0,95 | 0,99 | 0,01 | -168,23 | *NOD2* | 16 |
| ch.1.839062R | 0,87 | 0,78 | 0,97 | 0,01 | -333,21 | *RUNX3* | 1 |
| cg12688670 | 0,97 | 0,94 | 0,99 | 0,01 | -322,50 | *KIF22* | 16 |
| cg13066481 | 0,98 | 0,96 | 0,99 | 0,01 | -228,44 | *MYLK* | 3 |
| cg26166595 | 0,98 | 0,96 | 1,00 | 0,01 | -211,39 | *DKK3* | 11 |
| cg00863306 | 0,91 | 0,84 | 0,98 | 0,01 | -301,57 | *NANOS3* | 19 |
| cg04424621 | 0,94 | 0,89 | 0,99 | 0,01 | -249,37 | *HIST1H2BJ* | 6 |

| | | | | | | | |
|---|---|---|---|---|---|---|---|
| cg18993949 | 0,98 | 0,96 | 1,00 | 0,01 | -263,76 | *CC2D2A* | 4 |
| cg21572722 | 1,03 | 1,01 | 1,05 | 0,01 | 333,99 | *ELOVL2* | 6 |
| cg06142740 | 0,98 | 0,96 | 1,00 | 0,01 | -300,47 | *PPP1CB* | 2 |
| cg07368443 | 0,98 | 0,96 | 1,00 | 0,01 | -295,40 | *PLK3* | 1 |
| cg13807549 | 0,98 | 0,96 | 1,00 | 0,01 | -220,42 | | 9 |
| cg05045027 | 0,98 | 0,97 | 1,00 | 0,01 | -226,11 | | 11 |
| cg13428009 | 0,85 | 0,75 | 0,97 | 0,01 | -371,17 | | 9 |
| cg15416179 | 0,80 | 0,68 | 0,96 | 0,01 | -310,10 | *MAP2K3* | 17 |
| cg26276120 | 0,96 | 0,93 | 0,99 | 0,01 | -312,80 | *TPI1* | 12 |
| cg25371036 | 0,98 | 0,96 | 1,00 | 0,02 | -231,10 | *AMOTL1* | 11 |
| cg16363586 | 0,95 | 0,91 | 0,99 | 0,02 | -241,65 | *BST2* | 19 |
| cg19344626 | 0,99 | 0,97 | 1,00 | 0,02 | -160,82 | *NWD1* | 19 |
| cg20964856 | 0,98 | 0,96 | 1,00 | 0,02 | -271,65 | *HNRNPUL1;AXL* | 19 |
| cg08913523 | 0,97 | 0,94 | 0,99 | 0,02 | -189,17 | | 8 |
| cg02010481 | 0,97 | 0,94 | 0,99 | 0,02 | -158,13 | *JAZF1* | 7 |
| cg06661266 | 0,99 | 0,98 | 1,00 | 0,02 | -200,46 | | 3 |
| cg11807280 | 0,99 | 0,98 | 1,00 | 0,02 | -120,85 | | 2 |
| cg18786171 | 0,98 | 0,97 | 1,00 | 0,02 | -248,96 | *ADK* | 10 |
| ch.1.171672612F | 0,77 | 0,62 | 0,96 | 0,02 | -347,67 | | 1 |
| cg02610723 | 0,96 | 0,93 | 1,00 | 0,03 | -330,23 | *FAM38A* | 16 |
| cg03206537 | 0,97 | 0,94 | 1,00 | 0,03 | -199,68 | *CTSA* | 20 |
| cg16677191 | 0,97 | 0,94 | 1,00 | 0,03 | -191,76 | *GLRX* | 5 |
| cg18215449 | 0,99 | 0,98 | 1,00 | 0,03 | -177,35 | | 12 |
| cg09124496 | 0,99 | 0,98 | 1,00 | 0,03 | -176,00 | *LOC285954;INHBA* | 7 |
| cg24847230 | 0,98 | 0,96 | 1,00 | 0,03 | -345,09 | *UBE2Z* | 17 |
| cg05619598 | 0,98 | 0,97 | 1,00 | 0,03 | -197,92 | | 19 |
| cg06285727 | 0,95 | 0,91 | 1,00 | 0,03 | -213,65 | *ATG16L2* | 11 |
| cg26969888 | 0,98 | 0,96 | 1,00 | 0,03 | -201,74 | *PODNL1;DCAF15* | 19 |
| ch.6.33611621F | 0,63 | 0,42 | 0,96 | 0,03 | -328,25 | | 6 |
| cg00103778 | 0,98 | 0,96 | 1,00 | 0,03 | -217,65 | | 20 |
| cg14977938 | 1,02 | 1,00 | 1,05 | 0,03 | -258,46 | *ZFYVE21* | 14 |
| cg07843120 | 0,93 | 0,88 | 0,99 | 0,03 | -264,09 | *GPI* | 19 |
| ch.15.67797584R | 0,92 | 0,84 | 0,99 | 0,04 | -236,06 | | 15 |
| cg19056004 | 0,99 | 0,97 | 1,00 | 0,04 | -227,46 | *LRRC23;ENO2* | 12 |
| cg26543112 | 0,97 | 0,95 | 1,00 | 0,04 | -234,51 | | 6 |
| cg08957484 | 0,99 | 0,97 | 1,00 | 0,04 | 198,18 | *CCNI2* | 5 |
| cg16810343 | 0,98 | 0,97 | 1,00 | 0,04 | -244,57 | *SRRM3* | 7 |
| cg00753885 | 0,97 | 0,94 | 1,00 | 0,04 | -352,74 | | 12 |
| cg08301612 | 0,98 | 0,96 | 1,00 | 0,04 | -175,99 | *HEXIM1* | 17 |
| cg01981760 | 0,95 | 0,90 | 1,00 | 0,04 | -423,95 | *FTO;RPGRIP1L* | 16 |
| cg19784428 | 0,99 | 0,97 | 1,00 | 0,04 | -185,42 | *NWD1* | 19 |
| cg23744638 | 0,99 | 0,98 | 1,00 | 0,04 | -177,22 | | 11 |
| cg12179661 | 0,98 | 0,97 | 1,00 | 0,05 | -269,98 | *ENTPD8* | 9 |
| cg15034393 | 0,98 | 0,97 | 1,00 | 0,05 | -150,56 | | 3 |
| cg25424279 | 0,98 | 0,96 | 1,00 | 0,05 | -260,30 | | 11 |
| cg02797271 | 0,93 | 0,86 | 1,00 | 0,05 | -311,15 | *GPR132* | 14 |
| cg26210267 | 0,97 | 0,94 | 1,00 | 0,05 | -303,00 | *ATP5I* | 4 |
